# RegulaTome: a corpus of typed, directed, and signed relations between biomedical entities in the scientific literature

**DOI:** 10.1101/2024.04.30.591824

**Authors:** Katerina Nastou, Farrokh Mehryary, Tomoko Ohta, Jouni Luoma, Sampo Pyysalo, Lars Juhl Jensen

## Abstract

**Motivation:** In the field of biomedical text mining, the ability to extract relations from literature is crucial for advancing both theoretical research and practical applications. There is a notable shortage of corpora designed to enhance the extraction of multiple types of relations, particularly focusing on proteins and protein-containing entities such as complexes and families, as well as chemicals.

**Results:** In this work we present RegulaTome, a corpus that overcomes the limitations of several existing biomedical relation extraction (RE) corpora, many of which concentrate on single-type relations at the sentence level. RegulaTome stands out by offering 16,962 relations annotated in over 2,500 documents, making it the most extensive dataset of its kind to date. This corpus is specifically designed to cover a broader spectrum of over 40 relation types beyond those traditionally explored, setting a new benchmark in the complexity and depth of biomedical RE tasks. Our corpus both broadens the scope of detected relations and allows for achieving noteworthy accuracy in RE. A Transformer-based model trained on this corpus has demonstrated a promising F1-score (66.6%) for a task of this complexity, underscoring the effectiveness of our approach in accurately identifying and categorizing a wide array of biological relations. This achievement highlights RegulaTome’s potential to significantly contribute to the development of more sophisticated, efficient, and accurate RE systems to tackle biomedical tasks. Finally, a run of the trained relation extraction system on all PubMed abstracts and PMC Open Access full-text documents resulted in over 18 million relations, extracted from the entire biomedical literature.

**Availability:** The corpus and all introduced resources are openly accessible via Zenodo (https://zenodo.org/doi/10.5281/zenodo.10808330) and GitHub (https://github.com/farmeh/RegulaTome_extraction).

## Introduction

In the rapidly evolving field of Biomedical Natural Language Processing (BioNLP) and text mining, new technologies allow researchers to discover relations between biomedical entities. Relation Extraction (RE) is a critical task that enables the identification of relations among named entities (NE) such as genes, chemicals, and diseases. This process is essential for transforming unstructured text into structured data that can be used in both biological [Szklarczyk et al., 2023] and medical [Lee et al., 2016] applications.

The effectiveness of modern RE methodologies, particularly those leveraging the capabilities of pre-trained transformer models tailored for the biomedical domain [Lee et al., 2019, Lewis et al., 2020], hinges on the size, quality, and scope of manually annotated corpora used for model fine-tuning. These corpora serve as training and evaluation resources, guiding the development of methods capable of accurate information extraction. However, a majority of currently available corpora for RE are constrained by either focusing on relations at the sentence level [Bunescu et al., 2005, Pyysalo et al., 2007, Miranda-Escalada et al., 2023, Herrero-Zazo et al., 2013] and/or relations between two types of entities only (e.g. gene-disease) [Bunescu et al., 2005, Krallinger et al., 2008, Herrero-Zazo et al., 2013, Li et al., 2016]. Such constraints limit the number of relations that can be effectively extracted from the literature.

Recognizing these limitations, the BioNLP community has begun to shift its focus towards the development of more comprehensive corpora that extend beyond the sentence level to encompass document-level annotations [Doughty et al., 2011, Li et al., 2016, Su et al., 2021, Luo et al., 2022]. Standing out amongst them, the recent BioRED corpus [Luo et al., 2022], also tackles the issue of constrained scope, by having a broader coverage of 8 different relation types amongst disease, gene, variant, and chemical entities. While there are event annotation corpora that primarily concentrate on proteins and related entities and offer many document-level event annotations ([Kim et al., 2009, Ohta et al., 2010, Pyysalo et al., 2012]), a noticeable absence remains in an RE corpus with the same properties.

In this work, we introduce RegulaTome, a corpus comprising 2,521 documents with 16,962 document-level annotations, encompassing more than 40 types of relations — aligning with Gene Ontology [Aleksander et al., 2023, Ashburner et al., 2000] — between 54,951 entities belonging in four different entity types. This corpus is specifically designed to illuminate the complex web of interactions between proteins and protein-containing entities, providing an invaluable resource for advancing the state of RE in the biomedical field. Using this corpus, we have trained a transformer-based model with commendable results on relation extraction (F1-score=66.6%) for such a difficult task. To achieve this, we have developed an RE system capable of multi-label extraction of these directed, typed, and signed relations from the entire biomedical literature. This work fills a critical gap in biomedical relation extraction, offering a corpus and a system that allows the investigation of the complex interplay between proteins, protein-containing entities, and chemicals, which is foundational to understanding biological processes and disease mechanisms.

## Materials and methods

### The RegulaTome corpus

#### Targeted relation types

As mentioned above the aim of this work was to allow the extraction of directed, typed, and signed relations for proteins, chemicals, protein-containing complexes, and protein families from the literature. As many relation types between biomedical entities can fulfill these criteria, in this section we provide a list of the relation types that we have decided to annotate. We have mapped and structured the relation type space on the *Biological Process* sub-ontology of Gene Ontology (GO) [Aleksander et al., 2023, Ashburner et al., 2000], a community-standard framework. The full list of targeted relation types, the GO term corresponding to each of them, and their direct parent in our sub-ontology of relations is given in *Supplementary Section 1*. Figure 1 A shows an overview of the relationship tree, while Figure 1 B shows the relation representations within RegulaTome (for more details on the latter please refer to section *“Named Entity and Relation Annotation”*).

**Fig. 1.**
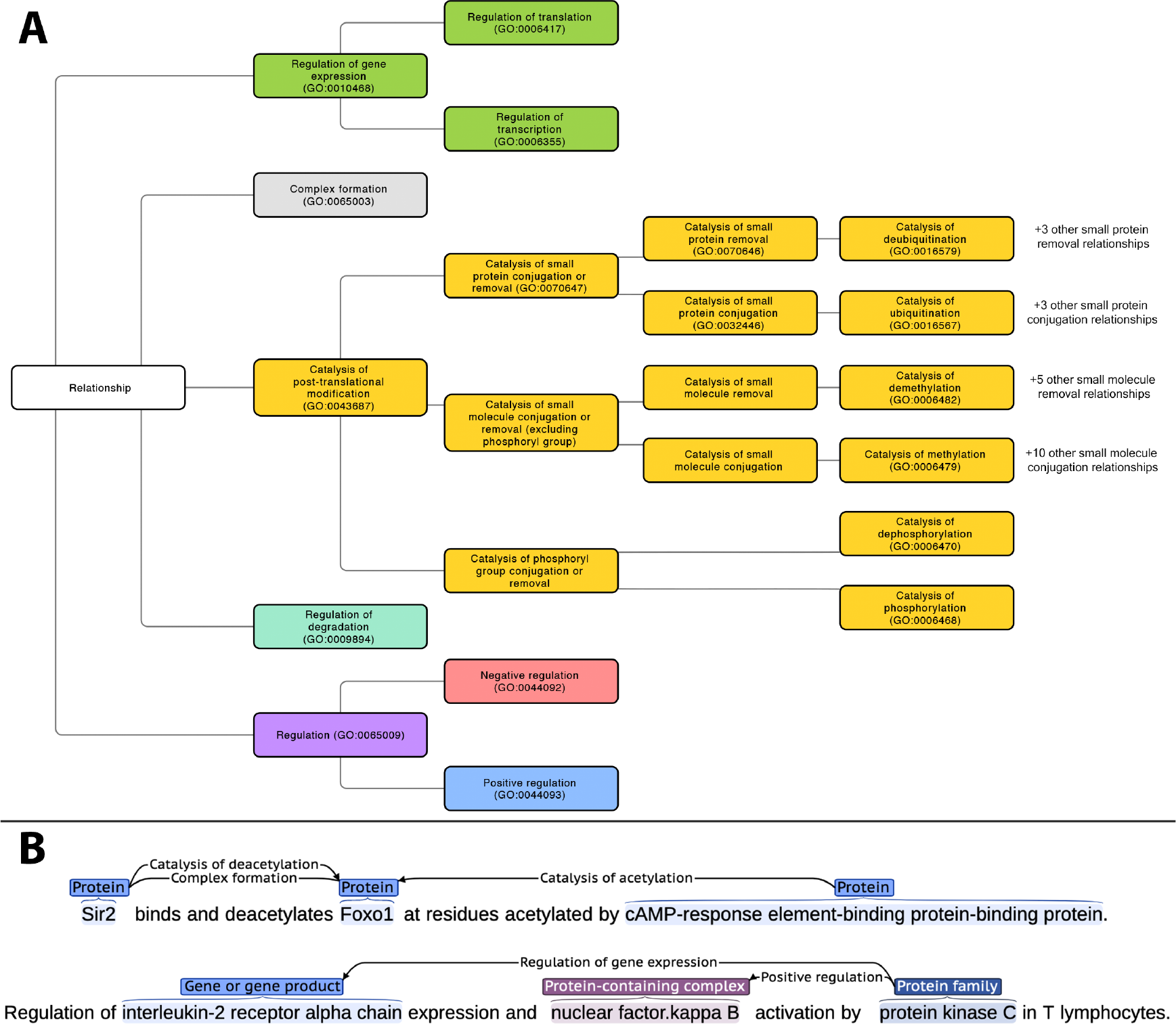
A. Targeted relation types in RegulaTome and their relationship to each other. There are 43 relation types annotated in RegulaTome, mapped to the `Biological Process` sub-ontology of GO. If the term is right below the root of our relation ontology, “Relationship” is used as a parent. Since GO lacks terms to collectively catalog all `Catalysis of small molecule conjugation or removal` processes, we have decided to group `Catalysis of phosphoryl group conjugation or removal` relations (i.e. `Catalysis of phosphorylation` and `Catalysis of dephosphorylation`) separately from the other `Catalysis of small molecule conjugation/removal` relations, both because of their biological significance, and — most importantly — because we observed that these are discussed differently in the biomedical literature. **B. Illustration of relation representations in RegulaTome**. (Top) multiple relations between a `Protein` (“Sir2”) and another `Protein` (“Foxo1”) participant are shown. Specifically, an undirected `Complex formation` relation and a directed `Catalysis of deacetylation` relation denoted with a left-to-right arrow, originating from “Sir2” with “Foxo1” as the target. There is one more directed relation that has “Foxo1” as the target, this time originating from another `Protein` participant (“cAMP-response element-binding protein-binding protein”) and in the opposite direction denoted by a right-to-left arrow. (Bottom) Relationships can arise between all entity types annotated. In this example, two directed relations (`Regulation of gene expression`, `Positive Regulation`) originate from a `Family` (“protein kinase C”) participant and target a `Protein` participant and a `Complex` (“nuclear factor.kappa B”).

#### Document selection for corpus annotation

The document selection process for the RegulaTome corpus consists of four steps:

##### 1. ComplexTome corpus [Mehryary et al., 2023]

this corpus consists mostly of documents focusing on physical protein interactions. This corpus includes 137 abstracts with complex formation events from BioNLP ST 2009 datasets [Kim et al., 2009] and 450 abstracts and 400 paragraphs from full-text used as evidence to support interactions in the BioGRID [Oughtred et al., 2021], IntAct [Orchard et al., 2014] and MINT [Licata et al., 2012] interaction databases. Moreover, this corpus contains 300 abstracts used for pathway annotation in the Reactome pathway knowledgebase [Gillespie et al., 2022] where regulatory relations are expected to be found in high prevalence. More details on the document selection of this corpus can be found in Mehryary et al. [2023]. We reannotated all documents of ComplexTome to include all 43 relation types mentioned in *Supplementary Section 1*.

##### 2. Post-Translational Modifications Event Extraction corpus

Out of the 388 abstracts in this corpus originally annotated for the BioNLP 2010 workshop [Ohta et al., 2010], we selected 234 based on each document containing at least one Post-Translational Modification (PTM) event. We ignored the existing event annotations and completely re-annotated the documents with relevant relations.

##### 3. PTM triage set

A pool of 5,548 publications from the Reactome database was generated by selecting those used to annotate pathways with at least one modification enzyme participant by the database curators [Gillespie et al., 2022]. We then went through the abstracts of these publications to select 500 of them, if there was at least one catalysis of PTM relation of interest therein. The selection was done incrementally with sets of 100 documents at a time and focusing on PTM relations with a lower number of total annotations every round to increase support for more relation types. *Supplementary Section 2* provides more details on the selection process.

##### 4. Reactome full-text excerpts set

A set of paragraphs from full-text articles used as evidence for pathway annotation in Reactome were selected if (a) they contained between 50 and 500 words — thus excluding documents with only titles or excessively lengthy paragraphs — (b) the number of unique tagged named entities within these paragraphs —disregarding case sensitivity — exceeded three entities, and (c) at least 30% of the entity mentions in each selected paragraph were forms not previously encountered in the documents of the corpus — to increase diversity. If a paragraph was chosen from a document for which its abstract is already included in our dataset, both the paragraph and the abstract were later assigned to the same subset (training, development, or test). There were 61,973 paragraphs from 21,941 papers fulfilling the criteria above, out of which we selected 500 for annotation. Similarly to the “PTM triage set”, selection was done in batches of 100 documents at a time. After initial observations, we tried to focus our annotations on specific sections of scientific papers, where most relations were expected to be mentioned. *Supplementary Section 3* provides more details on the process.

#### Named entity and relation annotation

There are four NE types in this corpus: Gene or gene products (`Protein` hereafter), chemicals (`Chemical`), protein-containing complexes (`Complex` hereafter), and protein families (`Family` hereafter). To annotate `Complex` entities we have used the definition of the homonymous term from GO (GO:0032991). As for `Family` entities, we have only annotated entities that are evolutionarily related, using InterPro [Paysan-Lafosse et al., 2023] as the main reference resource. Equivalent names of the same entities are systematically annotated to ensure evaluation accuracy [Kim et al., 2009].

In RegulaTome we identified explicit mentions of over 40 different relation types (*Supplementary Section 1*) and annotated those as either undirected (`Complex formation`) or directed (all other types) relations. Each candidate entity pair could receive multiple labels without any restrictions, and directed relations between the same entities could be bi-directional. Two examples of relation representations are shown in Figure 1 B.

Two experts in the field carried out the relation annotations for RegulaTome. An Inter-Annotator Agreement (IAA) analysis was performed to set uniform annotation standards and preserve the quality of annotations. This involved independently annotating a collection of the same abstracts in seven rounds. Four rounds of independent annotations were conducted on 80 documents from *ComplexTome* to establish the original annotation guidelines that acted as a guide to ensure annotators had a common understanding of them, aiding in the upkeep of high-quality annotations. Three additional rounds of IAA were conducted on 90 documents from the *Post-Translational Modifications Event Extraction corpus*. This resulted in a set of updated guidelines for the entire corpus and a re-annotation of all documents based on the updated set of rules. After each round, we measured the F1-score for IAA to evaluate the consistency of the annotations and the quality of the corpus. For detailed information on the annotation guidelines used to annotate NEs and relations in RegulaTome, we direct readers to the annotation documentation (available via Zenodo). The BRAT Rapid Annotation Tool [Stenetorp et al., 2012] was used for the annotation of all documents in RegulaTome.

#### Relation extraction system

We have extended the transformer-based relation extraction system we had previously developed for binary relation extraction [Mehryary et al., 2023] and developed our current system which is capable of extracting the relation types presented in *Supplementary Section 1*, between all NE types mentioned above. The task of RE is cast as a multi-label classification problem, where the goal is to predict if a pair of candidate NEs in the input text has one, several, or no stated relations. For the undirected `Complex formation` relation, there is only one dimension in the decision layer of the neural network, whereas for *each* directed relation, there are two dimensions, from the first occurring entity to the second occurring entity (i.e. left-to-right) and from the second occurring entity to the first (i.e. right-to-left).

Similarly to the binary classification system upon which we built, our current system utilizes an architecture featuring a pre-trained transformer encoder and a decision layer with a sigmoid activation function. The system can utilize pre-trained language models available in the Hugging Face repository, accepts training, validation, and prediction data in BRAT standoff and a custom JSON format, and supports extensive hyper-parameters (including maximum sequence length (MSL), learning rate, number of training epochs, and batch size). Evaluation metrics are calculated after each training epoch for hyper-parameter tuning. The system does not use any early stopping rule but is trained for a specified number of epochs and choosing the model weights that have yielded the highest F1-score.

The documents in our corpus, as is typical for biomedical documents, contain multiple candidate NE pairs and can be long, exceeding the maximum token capacity of transformer models. To clarify for the classifier which two candidate NEs are being considered for label prediction at a time, we encode these entities within the document using a masking approach, employing the model’s “unused” tokens for this purpose. We then tokenize the text and consider a window (a text snippet) around and including the two NEs (based on the MSL) and insert a [CLS] token at the beginning to signify the start of the snippet and a [SEP] token at the end of the input. Then for each candidate NE pair, we check if the masked, tokenized snippet that constitutes this pair, does not exceed the MSL we have specified and then create a machine-learning example for that pair. This example could be assigned one, multiple, or no labels for training, or remain unlabeled for prediction. Since we do not employ any sentence boundary detection, we can train with and predict cross-sentence relations on a document level. Moreover, relying on a window that can always be fed to the transformer encoder, allows us to effortlessly deal with long texts. If a candidate NE pair (i.e. a machine learning example) does not fit into the specified window size, it will be excluded in training and prediction, and penalized in the evaluation of development and test sets (if the two NEs have any relations between them).

For more details on the implementation and the strategy for pre-processing, input representation, and example generation, please refer to Mehryary et al. [2023].

#### Experimental setup

We performed a document-based split of RegulaTome into separate training, development, and test sets for our experiments. We use grid search to find the optimal values of hyper-parameters. To minimize the impact of initial random weights in neural network models on evaluation metrics [Mehryary et al., 2016], we repeat each *experiment* four times and compare different experiments based on the average and standard deviation of the F1-scores. Each experiment consists of training a relation extraction system (i.e. a neural network model) with the exact set of hyper-parameters but different initial random weights on the training set and evaluating the model on the development set. The hyper-parameter set which yields the highest average F1-score is chosen as the optimal, and the model with the highest F1-score in that experiment is selected for predicting the held-out test set and for large-scale execution of the relation extraction system on biomedical literature. Therefore, the test set is used only once for evaluating our best model.

## Results and discussion

### Corpus statistics

RegulaTome is a corpus of high quality that contains 2,521 documents with one paragraph each (1,621 abstracts and 900 paragraphs from full-text articles) consisting of approximately 612,000 words. The corpus quality was assessed through seven rounds of Inter-Annotator Agreement (IAA), which resulted in a final F1-score of 91% for IAA of all relation types. RegulaTome includes a total of 16,962 relations, with 6,463 of them being `Complex formation` (∼38%), followed by 2,294`Regulation` relations, 2,131 `Positive Regulation` and 1,920 `Negative Regulation`. *Supplementary Section 4* has annotation statistics for all relation types and *Supplementary Section 5* has the distribution of relations in the training, development, and test sets. RegulaTome offers a vast and varied set of relations for training neural network models for multi-label relation extraction. More than 95% of these relations occur within sentences and the rest span across sentences. The corpus also features a significant number of NEs, with 38,931 `Protein`, 4,703 `Chemical`, 3,839 `Complex`, and 7,478 `Family` NEs present, summing to 54,951 entities for all entity types.

### Relation extraction system evaluation

We used an extended grid search to find the optimal values of hyper-parameters on the development set of the RegulaTome corpus. Our best result was achieved using the `RoBERTa-large-PM-M3-Voc` model [Lewis et al., 2020] and the following set of hyper-parameters: *MSL=128, learning rate=4e-6, training epochs=26, batch size=16*.

Our best experiment achieved an average precision of 68.9%, an average recall of 67.0%, and an average F1-score of 67.9% on the RegulaTome development set. The four models we used in this experiment and the evaluation scores we measured on the development set are shown in Table 1.

**Table 1.** Performance of the best experiment on the RegulaTome development set. The best model (highlighted in gray) is used to perform a run on the held-out test set and for a large-scale run on the entire biomedical scientific literature

|  | Precision | Recall | F1-score |
| --- | --- | --- | --- |
| model-1 | 69.1 | 67.4 | 68.3 |
| model-2 | 68.3 | 66.3 | 67.3 |
| model-3 | 69.7 | 66.8 | 68.2 |
| model-4 | 68.6 | 67.3 | 67.9 |
| average | 68.9 | 67.0 | 67.9 |
| std | 0.61 | 0.51 | 0.45 |

The best model presented in Table 1 (model-1) achieved 66.6% F1-score (67.7% precision, 65.5% recall) on the RegulaTome held-out test set.

In *Supplementary Section 6*, evaluation metrics on the test set are presented on a per relation label basis. `Complex formation` — the label with the highest level of support — is, unsurprisingly, among the relations where the model achieves its best performance (F1-score=78.8%). Performance varies significantly for `Catalysis of posttranslational modification` relations, with F1-scores varying from 85.7% for `Catalysis of deubiquitination` to 0% for e.g. `Other catalysis of small protein removal`. Results in these cases seem to be directly affected by the level of support per label (*Supplementary Section 4*), with labels with a higher level of support like `Catalysis of ubiquitination`, `Catalysis of phosphorylation`, `Catalysis of dephosphorylation`, and `Catalysis of methylation`, having F1-scores around 70%. `Regulation`-related labels seem to be the most difficult to predict, a result consistent with the literature on similar tasks [Miranda-Escalada et al., 2023]. Relationship sign assignment seems to be easier than the general class prediction, with `Positive Regulation` and `Negative Regulation` having F1-scores above 62%, while `Regulation`, despite its high level of support, achieving an F1-score of only 49.3%. Moreover, `Regulation of transcription` seems easier to predict than `Regulation of gene expression`, but this could again be explained by the fact that the level of support for `Regulation of transcription` is double that of `Regulation of gene expression` (*Supplementary Section 4*).

In the next sections, we perform a manual error analysis and a semi-automated label confusion analysis, which allows us to look deeper into these results.

### Manual error analysis

We have selected 20% of documents in the test set and manually analyzed and categorized the errors generated by the best-performing relation extraction model on these documents. An overview of these errors is shown in Table 2, while a case-by-case analysis is provided in *Supplementary Section 7*.

**Table 2.** Manual error analysis on 20% of documents in the RegulaTome test set.

| Error Type | Count |  |  |
| --- | --- | --- | --- |
|  | FP | FN | Total |
| Ambiguous keyword | 65 | 35 | 100 |
| Rare keyword | 0 | 57 | 57 |
| Co-reference resolution | 31 | 29 | 60 |
| Convoluted text excerpt | 61 | 55 | 116 |
| Model error | 17 | 32 | 49 |
| Annotation error | 26 | 15 | 41 |
| Total | 200 | 223 | 423 |

\*FN: false negative, FP: false positive

From the error categories presented in Table 2 the main sources of errors appear to be “ambiguous keyword” and “convoluted text excerpt”, with over half of the errors being a result of these. The first category encapsulates instances where ambiguous words, like “target”, can denote either a regulatory (e.g. *The promoter of the CD19 gene is a* ***target*** *for BSAP*) or a catalytic (e.g. *Tea1 is a substrate* ***target*** *of Shk1*) relation, and result in model confusion. The second most common category (“convoluted text excerpt”) encompasses text segments with complex syntax, including intricate sentences and cross-sentence relations, which are inherently difficult to annotate and subsequently predict. A category closely related to that is “co-reference resolution”, where the syntactical structure makes it especially difficult for the model to assess which subject a given relation pertains to and results in both False Positives (FPs) and False Negatives (FNs). The “rare keyword” category results only in FNs as a consequence of words or phrases rarely found in scientific texts (e.g. *protection from inhibition* or *non-covalent association*), which are recognized and correctly annotated by biology experts, but do not result in enough examples for the model to train on to have a chance to detect them during prediction.

There are two more categories — with lower numbers of errors — which are inherently different than the rest of the categories presented above. “Model error” refers to cases where there are clear keywords to denote relations, and where there were no clear explanations as to why these have not been correctly predicted by the model. On the other hand, “annotation error” refers to cases in which annotators have inaccurately labeled or not labeled relations, frequently as a result of text ambiguity, which would require correction in the corpus.

### Label confusion analysis

Next, we have categorized the errors based on the confusion of relation labels (*Supplementary Sections 8* and *9*). Overall, the vast majority of all FPs (81%) are cases where relations are predicted and there should be no relation of any type according to our manual annotations (*Supplementary Section 8*, bold & italics). Similarly, 82% of FNs are relations that were completely missed (*Supplementary Section 9*, bold & italics) and are not a result of confusion between labels predicted by the model. For a full categorization of each FP and FN in the RegulaTome test set in terms of label confusion, please refer to the Supplementary Table (*Error analysis full results*) available via Zenodo.

For the remaining errors, some label confusion categories are less severe than others. Specifically, 10% of all errors in the test set (126 out of 1048 FPs and 118 out of 1160 FNs) — i.e. half of the remaining errors — have to do with confusion among closely related labels (*Supplementary Sections 8* and *9*). For example, in the `Regulation of gene expression` branch (Figure 1) either a too-specific label (i.e. `Regulation of transcription` instead of `Regulation of gene expression`) or a too-broad label (i.e. `Regulation of gene expression` instead of `Regulation of transcription`) was predicted. If all confusion within the `Regulation of gene expression` branch was ignored, i.e. if all confusion between `Regulation of transcription`, `Regulation of translation`, and `Regulation of gene expression` labels is counted as True Positives (TPs) instead of FPs and FNs, the average F1-score for the `Regulation of gene expression` branch increases to 68.8%, which is 9% better than `Regulation of transcription` and 15% better than `Regulation of gene expression` (*Supplementary Section 6*). Similarly, if all confusion within the `Catalysis of posttranslational modification` branch is ignored, the average F1-score for `Catalysis of posttranslational modification` increases to 70.6%, which is better than the F1-scores for 18 of the 22 relation types within that branch (*Supplementary Section 6*).

### Error analysis of direction and sign

The directed relations that can be mined from literature using our model can provide important information for the analysis of regulatory networks. In this use case, relations are viewed as edges, and what matters most is to have the correct edges, with the right direction, and ideally the right sign (i.e. `Positive Regulation` or `Negative Regulation`). To evaluate the usefulness of our model’s predictions for this purpose, we categorized label confusion errors in six categories, considering only directed predictions and annotations (i.e. the presence or absence of predicted or annotated `Complex formation` has no impact), namely cases where the model

1. failed to assign a directed interaction, where there should be one
2. assigned a directed interaction, where there should be none
3. assigned a directed interaction, but the direction is wrong
4. failed to assign a sign (positive or negative), where there should be one
5. assigned a sign, where there should be none
6. assigned a sign, but the sign is wrong

We found 1,394 edges with correctly assigned directions and 737, 620, and 5 errors from the first three categories, respectively. While the network that would be produced is somewhat incomplete — missing 737 interactions — its precision would be 70% in terms of connecting the right entities with an edge pointing the right way. It should be noted that in reality, the precision would be even higher since some of the relations counted as FPs are annotation errors in the corpus. Of the correctly detected directed edges, 539 of them furthermore have the correct sign, while 31 are missing a sign (category 4), and 61 have a wrong sign (47 from category 5 and 14 from category 6). For the remaining 763 edges we correctly did not predict a sign. These results further showcase the potential of deep learning-based models trained on RegulaTome for downstream biomedical applications. For details on calculations presented in this section please refer to *Supplementary Section 10*.

### Large-scale execution for protein relations

We used the best model to extract relations from over 36 million PubMed abstracts (as of March 2024) and 6 million articles from the PMC BioC open access collection [Comeau et al., 2019] (as of November 2023). The Jensenlab tagger [Jensen, 2016] was used to obtain matches for `Protein` named entities with normalizations to Ensembl [Martin et al., 2022] identifiers, and the results were filtered to documents that contain at least two named entities and as a result at least one pair for prediction. 6,920,139 documents complied with this criterion (3,157,239 abstract and full-text and 3,762,900 abstract only) which were converted to BRAT standoff format and provided to the model for relation prediction. Predictions were produced for over 1.2 billion pairs, with ∼1.5% (18.4 million) having at least one “positive” label. A tab-delimited file with results from the large-scale run is provided through Zenodo.

## Conclusions

In this work, we introduced RegulaTome, a corpus aimed at enhancing biomedical RE, with a focus on proteins, protein-containing entities such as complexes and families, and chemicals. This work represents a significant advancement in the field of biomedical text mining, addressing a limitation of several existing RE corpora that mainly focus on single-type relations at the sentence level. RegulaTome distinguishes itself by its breadth, encompassing 2,521 documents with 16,962 relations between 54,951 entities. It is meticulously curated to include 43 types of relations, extending well beyond the scope traditionally covered in biomedical RE tasks, thereby establishing a new standard for complexity and depth in the field.

The effectiveness of RegulaTome is further demonstrated through the deployment of a transformer-based model, which has shown remarkable accuracy in RE, achieving an F1-score of 66.6% that underlines the corpus’s utility in accurately identifying and categorizing a diverse range of biological relations. This achievement showcases the corpus’s capacity to broaden the scope of detectable relations and its potential to significantly enhance the development of sophisticated, efficient, and accurate RE systems for biomedical applications. By providing RegulaTome to the scientific community, we aim to facilitate the advancement of biomedical RE systems both through theoretical research and practical applications in the field. Our work sets a new benchmark in biomedical text mining and opens up new avenues for exploring and validating a plethora of complex relations between biomedical entities.

## Supporting information

Supplementary Section

## Acknowledgements

We thank the CSC – IT Center for Science, Finland for generous computational resources.

