## Supplementary Section for "RegulaTome: a corpus of typed, directed, and signed relations between biomedical entities in the scientific literature"

### Supplementary Section 1: Relationship types in RegulaTome

| Relationship Type | GO ID | Parent Term |
|---|---|---|
| Complex formation | GO:0065003 | Relationship |
| Regulation | GO:0065009 | Relationship |
| Positive regulation | GO:0044093 | Regulation |
| Negative regulation | GO:0044092 | Regulation |
| Regulation of gene expression | GO:0010468 | Relationship |
| Regulation of transcription | GO:0006355 | Regulation of gene expression |
| Regulation of translation | GO:0006417 | Regulation of gene expression |
| Regulation of degradation | GO:0009894 | Relationship |
| Catalysis of post-translational modification | GO:0043687 | Relationship |
| Catalysis of small protein conjugation/removal | GO:0070647 | Catalysis of post-translational modification |
| Catalysis of small protein conjugation | GO:0032446 | Catalysis of small protein conjugation/removal |
| Catalysis of ubiquitination | GO:0016567 | Catalysis of small protein conjugation |
| Catalysis of SUMOylation | GO:0016925 | Catalysis of small protein conjugation |
| Catalysis of neddylation | GO:0045116 | Catalysis of small protein conjugation |
| Other catalysis of small protein conjugation | --- | Catalysis of small protein conjugation |
| Catalysis of small protein removal | GO:0070646 | Catalysis of small protein conjugation/removal |
| Catalysis of deubiquitination | GO:0016579 | Catalysis of small protein removal |
| Catalysis of deSUMOylation | GO:0016926 | Catalysis of small protein removal |
| Catalysis of deneddylation | GO:0000338 | Catalysis of small protein removal |
| Other catalysis of small protein removal | --- | Catalysis of small protein removal |
| Catalysis of phosphoryl group conjugation/removal | --- | Catalysis of post-translational modification |
| Catalysis of phosphorylation | GO:0006468 | Catalysis of phosphoryl group conjugation/removal |
| Catalysis of dephosphorylation | GO:0006470 | Catalysis of phosphoryl group conjugation/removal |
| Catalysis of small molecule conjugation/removal (excluding phosphoryl group) | --- | Catalysis of post-translational modification |
| Catalysis of small molecule conjugation | --- | Catalysis of small molecule conjugation/removal (excluding phosphoryl group) |
| Catalysis of methylation | GO:0006479 | Catalysis of small molecule conjugation |
| Catalysis of acylation | GO:0043543 | Catalysis of small molecule conjugation |

| Relationship Type | GO ID | Parent Term |
| --- | --- | --- |
| Catalysis of acetylation | GO:0006473 | Catalysis of acylation |
| Catalysis of palmitoylation | GO:0018345 | Catalysis of acylation |
| Catalysis of lipidation | GO:0006497 | Catalysis of small molecule conjugation |
| Catalysis of prenylation | GO:0018342 | Catalysis of lipidation |
| Catalysis of farnesylation | GO:0018343 | Catalysis of prenylation |
| Catalysis of geranylgeranylation | GO:0018344 | Catalysis of prenylation |
| Catalysis of ADP-ribosylation | GO:0070212 | Catalysis of small molecule conjugation |
| Catalysis of glycosylation | GO:0006486 | Catalysis of small molecule conjugation |
| Other catalysis of small molecule conjugation | --- | Catalysis of small molecule conjugation |
| Catalysis of small molecule removal | --- | Catalysis of small molecule conjugation/removal (excluding phosphoryl group) |
| Catalysis of demethylation | GO:0006482 | Catalysis of small molecule removal |
| Catalysis of deacylation | GO:0035601 | Catalysis of small molecule removal |
| Catalysis of deacetylation | GO:0006476 | Catalysis of deacylation |
| Catalysis of depalmitoylation | GO:0002084 | Catalysis of deacylation |
| Catalysis of deglycosylation | GO:0006517 | Catalysis of small molecule removal |
| Other catalysis of small molecule removal | --- | Catalysis of small molecule removal |

### Supplementary Section 2: Document selection for PTM triage set

Document selection for the construction of the PTM triage set involved five steps in total (`PTM triage 01` to `PTM triage 05`), all of which follow a similar logic, adapted based on relationship counts after each annotation round. The purpose of this expansion is to enrich the corpus in `Catalysis of protein modification` relationships, excluding `Catalysis of phosphorylation` for which we already had high support in the corpus.

#### PTM triage 01

To generate `PTM triage 01` we searched **UniProtKB** (accessed on January 24th, 2023) using the query `"(GO:0036211) AND (cc_catalytic_activity:*) NOT (GO:0016301) NOT (GO:0006468)"` to identify entries associated with the GO term `protein modification process` while excluding entries related to `protein kinase` and `protein phosphorylation`. This search yielded a total of 1,389,360 results, of which 12,522 were sourced from SwissProt and 1,376,838 from TrEMBL. Subsequently, files from the **Reactome** database were utilized to map the UniProt identifiers from the search above to Reactome reactions and to obtain associated PubMed identifiers (PMIDs) used to describe these reactions. Specifically, the `UniProt2Reactome_PE_Reactions.txt` file was used to map UniProt identifiers to Reactome pathway identifiers, and the `Reaction_PMIDs.txt` file provided information on research publications associated with these Reactome pathways. Reactions containing proteins from the initial list were extracted, resulting in 12,908 unique reactions, which linked to 5,548 unique PMIDs from research papers used to annotate them in Reactome. We then run an NER system on these abstracts, to identify those containing 2-40 entities (to be consistent with all selection steps in ComplexTome), which reduced the number of documents to 4,771. Afterwards we checked which of these documents we have already annotated (97 documents in total), and by removing those documents we are left with the final set of **4,674** documents. We converted these documents to JSON format, shuffled them to be in random order and set up a **prodigy** server to contain all the documents. This would allow us to manually inspect the documents to identify those that contain relationships of interest. This step was deemed necessary, as our previous effort to focus on documents containing `Catalysis of protein modification` relationships, by annotating documents in existing corpora (`Exhaustive PTM corpus`) resulted in very low density of the relationship types we were aiming for. For that reason we decided to "***triage***" the documents, by manually inspecting them and selecting those that contain the relationships of interest. After inspecting 850 documents we selected 100 of them,on the basis of mentioning a `catalysis of protein modification` relationship, besides `catalysis of phosphorylation`. These documents were later annotated and constitute the `PTM triage 01` set in our corpus.

#### PTM triage 02

For `PTM triage 02` we decided to focus on specific relationship types that were even more underrepresented in the corpus. We followed a similar document selection strategy, but this time we searched UniProt (accessed on March 4th, 2023) for entries that have a field of `catalytic activity` and a GO term for `Catalysis of methylation (GO:0006479)` or `Catalysis of acylation (GO:0043543)` or `Catalysis of lipidation (GO:0006497)` or `Catalysis of Glycosylation (GO:0006486)` or `Catalysis of Demethylation (GO:0006482)` or `Catalysis of Deacylation (GO:0035601)` which includes `Catalysis of deacetylation` and `Catalysis of depalmitoylation` or `Catalysis of Deglycosylation (GO:0006517)`. This search resulted in 380,014 entries; 4,094 from SwissProt and 375,920 from TrEMBL. We then searched Reactome for papers used to annotate reactions with these enzymes and filtered the resulting list to remove papers we already have in the corpus, or seen in the previous triage round and rejected. We also applied an extra filter on top to keep only papers with that contain at least on of the following lemmas (case insensitive) `acetyl|Acyl|add|ADPribosyl|ADP-ribosyl|attach|biotinyl|carbonyl|carboxyl|conjuga|Deacetyl|Deacyl|deconjug|Deglycosyl|deiminat|Demethyl|Deneddyl|Depalmitoyl|Dephosphoryl|DeSUMOyl|Deubiquitin|farnesyl|Gamma carboxyl|gamma-carboxyl|geranylgeranyl|GlcNAcyl|glycoform|Glycosyl|incorporat|isoprenyl|link|lipidmodif|lipidat|lipoyl|methyl|modif|monoacyl|monoprenyl|monoubiquitin|myristol|myristyl|N-glycosyl|neddyl|O-GlcNAcyl|O-glycosyl|O-link|O-Sulf|oxidat|Palmitoyl|Prenyl|remov|repalmitoylat|sulf|SUMO|ubiquitin`. This resulted in a set of 1,046 papers, which were again shuffled and added on prodigy for triage. After checking 428 of these abstracts, we selected 100 of them on the basis of containing the subset of `Catalysis of protein modification` relationships on which our UniProt search is also focused on, and created the `PTM triage 02` set.

#### PTM triage 03

Similarly to the creation we focused on specific relationship types during this round. Specifically, we focused on relationships that had a total support of less than 100 relationships in the annotated corpus up to that point and we searched Uniprot on March 28th, 2023 for entries that have a field of `"catalytic activity"` and a GO term for `Catalysis of ADP-ribosylation (GO:0070212)` or `Catalysis of SUMOylation (GO:0016925)` or `Catalysis of Acetylation (GO:0006473)` or `Catalysis of Neddylation (GO:0045116)` or `Catalysis of farnesylation (GO:0018343)` or `Catalysis of geranylgeranylation (GO:0018344)` or `Catalysis of Myristoylation (GO:0018377)` or `Catalysis of Glycosylation (GO:0006486)` or `Catalysis of Palmitoylation (GO:0018345)` or `Catalysis of Deubiquitination (GO:0016579)` or `Catalysis of DeSUMOylation (GO:0016926)` or `Catalysis of Deneddylation (GO:0000338)` or `Catalysis of`

`Dephosphorylation (GO:0006470)` or `Catalysis of Demethylation (GO:0006482)` or `Catalysis of Deacetylation (GO:0006476)` or `Catalysis of Depalmitoylation (GO:0002084)` or `Catalysis of Deglycosylation (GO:0006517)`. This search resulted in 431,963 entries; 4,473 from SwissProt and 427,490 from TrEMBL. We applied the exact same strategy to select documents from Reactome as for `PTM triage 02`, and ended up with a set of 1,229 papers, out of which 100 were selected if they contained any of the relationships mentioned above.

#### PTM triage 04

Once again the focus was on specific relationship types. In this selection round we searched Uniprot on May 8th, 2023 for entries with a field of `"catalytic activity"` and a GO term for `Catalysis of small protein conjugation (GO:0032446)` or `Catalysis of small protein removal (GO:0070646)` or `Catalysis of Dephosphorylation (GO:0006470)` or `Catalysis of Demethylation (GO:0006482)` or `Catalysis of Deacetylation (GO:0006476)` or `Catalysis of Depalmitoylation (GO:0002084)` or `Catalysis of Deglycosylation (GO:0006517)`. This search resulted in 487,244 entries; 4,478 from SwissProt and 482,766 from TrEMBL. We applied the exact same strategy to select documents from Reactome as for `PTM triage 02` and `03`, and once more selected 100 documents out of 900 if they contained any of the relationships mentioned above to generate `PTM triage 04`.

#### PTM triage 05

For `PTM triage 05` we continued using the same documents as for PTM triage 04, but ignored documents mentioning `Catalysis of small protein removal (GO:0070646)` during triage. We selected 100 more documents this way and thus concluded all the selection steps for the `PTM triage` set.

#### Supplementary Section 3: Document selection for Reactome full-text excerpts set

For ComplexTome we had already selected 300 abstracts extracted from 21,941 papers used for pathway annotation in the Reactome pathway knowledgebase. We used the same pool of papers to select 500 full-text paragraphs for the `Reactome full-text excerpts` set. We split these documents in paragraphs and ended up with a set of 384,696 total paragraphs. Then we used the codebase presented in the S1000 paper (*Luoma, et al, 2023*) to train a transformer-based model for NER of all named entity types annotated in the RegulaTome corpus, i.e. `Protein`, `Complex`, `Protein Family` and `Chemical`. We used the same document split we already have to generate a training, development and test set, and then used the named entity annotations to train a transformer-based model for multi-class classification of named entities. We used the best model based on a `RoBERTa-large-PM-M3-Voc` model and then predicted named entities in all the documents above. We shuffled the paragraphs and then used the following criteria to filter down the entire pool of paragraphs from which we would then select documents for each step of generating the `Reactome full-text excerpts` set:

- the paragraphs should contain at least three annotated textbound named entities of type `Protein`, `Complex` or `Protein Family`
- 30% of these names should have not been seen before in the corpus (so selection is done incrementally, one document at a time)
- the paragraph size should be between 50 and 500 words

After selecting documents with these criteria we were left with 61973 paragraphs from which we initially selected 200 documents to generate `Reactome full-text excerpts 01` and `Reactome full-text excerpts 02`. For the next 3 steps we created a custom ChatGPT (available through this link: https://chat.openai.com/g/g-s8Uk5qFoO-biomed-scholar) and instructed it to classify paragraphs from scientific papers into specific sections. We chose this option instead of mapping the paragraphs back to the original documents and identifying the sections, due to differences in how papers are structured in different journals, the lack of need of 100% accuracy in classification and the initial good classification results we were getting with the custom ChatGPT. For batches `Reactome full-text excerpts 03`, `Reactome full-text excerpts 04` and `Reactome full-text excerpts 05`, we filtered documents so that they were either from the Introduction, Results, Discussion or Conclusions sections (according to classification of our custom ChatGPT) and took extra care to manually remove any figure legends from `Reactome full-text excerpts 04` and `Reactome full-text excerpts 05` sets. We did that as we observed that paragraphs from sections other than those mentioned above mostly did not contain any of the relationships we annotate in this corpus, while we were annotating the first two batches of this set.

### Supplementary Section 4: Relationship statistics for RegulaTome

| Relationship type | Relationship count | Relationship type | Relationship count |
|---|---|---|---|
| Complex formation | 6463 | Catalysis of other small molecule conjugation or removal | 9 |
| Regulation | 2294 | Catalysis of small molecule conjugation | 0 |
| Positive regulation | 2131 | Catalysis of methylation | 259 |
| Negative regulation | 1920 | Catalysis of glycosylation | 43 |
| Regulation of gene expression | 521 | Catalysis of ADP-ribosylation | 31 |
| Regulation of transcription | 899 | Catalysis of acylation | 14 |
| Regulation of translation | 21 | Catalysis of acetylation | 129 |
| Regulation of degradation | 336 | Catalysis of palmitoylation | 66 |
| Catalysis of posttranslational modification | 169 | Catalysis of lipidation | 3 |
| Catalysis of small protein conjugation or removal | 5 | Catalysis of prenylation | 1 |
| Catalysis of small protein conjugation | 42 | Catalysis of farnesylation | 3 |
| Catalysis of ubiquitination | 474 | Catalysis of geranylgeranylation | 17 |
| Catalysis of SUMOylation | 86 | Other catalysis of small molecule conjugation | 33 |
| Catalysis of neddylation | 20 | Catalysis of small molecule removal | 2 |
| Other catalysis of small protein conjugation | 4 | Catalysis of demethylation | 110 |
| Catalysis of small protein removal | 1 | Catalysis of deglycosylation | 1 |
| Catalysis of deubiquitination | 79 | Catalysis of deacylation | 6 |
| Catalysis of deSUMOylation | 11 | Catalysis of deacetylation | 53 |
| Catalysis of deneddylation | 19 | Catalysis of depalmitoylation | 6 |
| Other catalysis of small protein removal | 3 | Other catalysis of small molecule removal | 16 |
| Catalysis of phosphoryl group conjugation or removal | 7 | | |
| Catalysis of phosphorylation | 442 | | |
| Catalysis of dephosphorylation | 213 | | |

### Supplementary Section 5: Relationship statistics in the training, development and test sets

| Relationship type | # Train | # Devel | # Test | % Train | % Devel | %Test |
|---|---|---|---|---|---|---|
| Complex formation | 3984 | 1258 | 1221 | 61.6% | 19.5% | 18.9% |
| Regulation | 1330 | 480 | 484 | 58.0% | 20.9% | 21.1% |
| Positive regulation | 1272 | 415 | 444 | 59.7% | 19.5% | 20.8% |
| Negative regulation | 1113 | 390 | 417 | 58.0% | 20.3% | 21.7% |
| Regulation of gene expression | 319 | 105 | 97 | 61.2% | 20.2% | 18.6% |
| Regulation of transcription | 559 | 168 | 172 | 62.2% | 18.7% | 19.1% |
| Regulation of translation | 8 | 6 | 7 | 38.1% | 28.6% | 33.3% |
| Regulation of degradation | 201 | 86 | 49 | 59.8% | 25.6% | 14.6% |
| Catalysis of posttranslational modification | 94 | 31 | 44 | 55.6% | 18.3% | 26.0% |
| Catalysis of small protein conjugation or removal | 3 | 0 | 2 | 60.0% | 0.0% | 40.0% |
| Catalysis of small protein conjugation | 29 | 8 | 5 | 69.0% | 19.0% | 11.9% |
| Catalysis of ubiquitination | 289 | 91 | 94 | 61.0% | 19.2% | 19.8% |
| Catalysis of SUMOylation | 57 | 15 | 14 | 66.3% | 17.4% | 16.3% |
| Catalysis of neddylation | 13 | 5 | 2 | 65.0% | 25.0% | 10.0% |
| Other catalysis of small protein conjugation | 3 | 0 | 1 | 75.0% | 0.0% | 25.0% |
| Catalysis of small protein removal | 1 | 0 | 0 | 100.0% | 0.0% | 0.0% |
| Catalysis of deubiquitination | 49 | 15 | 15 | 62.0% | 19.0% | 19.0% |
| Catalysis of deSUMOylation | 7 | 4 | 0 | 63.6% | 36.4% | 0.0% |
| Catalysis of deneddylation | 11 | 2 | 6 | 57.9% | 10.5% | 31.6% |
| Other catalysis of small protein removal | 2 | 0 | 1 | 66.7% | 0.0% | 33.3% |
| Catalysis of phosphoryl group conjugation or removal | 6 | 0 | 1 | 85.7% | 0.0% | 14.3% |
| Catalysis of phosphorylation | 252 | 91 | 99 | 57.0% | 20.6% | 22.4% |
| Catalysis of dephosphorylation | 126 | 42 | 45 | 59.2% | 19.7% | 21.1% |
| Catalysis of other small molecule conjugation or removal | 3 | 0 | 6 | 33.3% | 0.0% | 66.7% |

| Relationship type | # Train | # Devel | # Test | % Train | % Devel | %Test |
|---|---|---|---|---|---|---|
| Catalysis of methylation | 163 | 45 | 51 | 62.9% | 17.4% | 19.7% |
| Catalysis of glycosylation | 27 | 8 | 8 | 62.8% | 18.6% | 18.6% |
| Catalysis of ADP-ribosylation | 19 | 6 | 6 | 61.3% | 19.4% | 19.4% |
| Catalysis of acylation | 8 | 5 | 1 | 57.1% | 35.7% | 7.1% |
| Catalysis of acetylation | 74 | 26 | 29 | 57.4% | 20.2% | 22.5% |
| Catalysis of palmitoylation | 42 | 12 | 12 | 63.6% | 18.2% | 18.2% |
| Catalysis of lipidation | 3 | 0 | 0 | 100.0% | 0.0% | 0.0% |
| Catalysis of prenylation | 1 | 0 | 0 | 100.0% | 0.0% | 0.0% |
| Catalysis of farnesylation | 3 | 0 | 0 | 100.0% | 0.0% | 0.0% |
| Catalysis of geranylgeranylation | 17 | 0 | 0 | 100.0% | 0.0% | 0.0% |
| Other catalysis of small molecule conjugation | 27 | 2 | 4 | 81.8% | 6.1% | 12.1% |
| Catalysis of small molecule removal | 2 | 0 | 0 | 100.0% | 0.0% | 0.0% |
| Catalysis of demethylation | 73 | 19 | 18 | 66.4% | 17.3% | 16.4% |
| Catalysis of deglycosylation | 1 | 0 | 0 | 100.0% | 0.0% | 0.0% |
| Catalysis of deacylation | 3 | 3 | 0 | 50.0% | 50.0% | 0.0% |
| Catalysis of deacetylation | 35 | 9 | 9 | 66.0% | 17.0% | 17.0% |
| Catalysis of depalmitoylation | 5 | 1 | 0 | 83.3% | 16.7% | 0.0% |
| Other catalysis of small molecule removal | 2 | 14 | 0 | 12.5% | 87.5% | 0.0% |

### Supplementary Section 6: Evaluation statistics per class on the RegulaTome held-out test set

TP: True Positive, FN: False Negative, FP: False Positive

| Relationship | Precision | TP/TP+FP | Recall | TP/TP+FN | F-score |
|---|---|---|---|---|---|
| Catalysis of deubiquitination | 92.3% | 12/13 | 80.0% | 12/15 | 85.7% |
| Complex formation | 78.6% | 965/1227 | 79.0% | 965/1221 | 78.8% |
| Catalysis of demethylation | 69.6% | 16/23 | 88.9% | 16/18 | 78.0% |
| Catalysis of ubiquitination | 66.4% | 77/116 | 81.9% | 77/94 | 73.3% |
| Catalysis of phosphorylation | 76.5% | 65/85 | 66.3% | 65/98 | 71.0% |
| Catalysis of dephosphorylation | 68.1% | 32/47 | 71.1% | 32/45 | 69.6% |
| Catalysis of methylation | 70.0% | 35/50 | 68.6% | 35/51 | 69.3% |
| Catalysis of neddylation | 100.0% | 1/1 | 50.0% | 1/2 | 66.7% |
| Negative regulation | 64.8% | 263/406 | 63.2% | 263/416 | 64.0% |
| Positive regulation | 61.7% | 276/447 | 62.6% | 276/441 | 62.2% |
| Regulation of degradation | 51.4% | 36/70 | 73.5% | 36/49 | 60.5% |
| Regulation of transcription | 61.1% | 102/167 | 59.6% | 102/171 | 60.4% |
| Catalysis of acetylation | 60.7% | 17/28 | 58.6% | 17/29 | 59.6% |
| Catalysis of palmitoylation | 58.3% | 7/12 | 58.3% | 7/12 | 58.3% |
| Regulation of gene expression | 59.3% | 48/81 | 49.5% | 48/97 | 53.9% |
| Catalysis of SUMOylation | 58.3% | 7/12 | 50.0% | 7/14 | 53.8% |
| Catalysis of deacetylation | 45.5% | 5/11 | 55.6% | 5/9 | 50.0% |
| Regulation | 53.0% | 223/421 | 46.1% | 223/484 | 49.3% |
| Catalysis of ADP-ribosylation | 50.0% | 2/4 | 33.3% | 2/6 | 40.0% |
| Other catalysis of small molecule conjugation | 50.0% | 1/2 | 25.0% | 1/4 | 33.3% |
| Catalysis of small protein conjugation | 50.0% | 1/2 | 20.0% | 1/5 | 28.6% |
| Catalysis of posttranslational modification | 37.5% | 6/16 | 13.6% | 6/44 | 20.0% |
| Catalysis of glycosylation | 20.0% | 1/5 | 12.5% | 1/8 | 15.4% |
| Catalysis of deneddylation | 0.0% | 0/0 | 0.0% | 0/6 | 0.0% |
| Catalysis of other small molecule conjugation or removal | 0.0% | 0/0 | 0.0% | 0/6 | 0.0% |
| Catalysis of phosphoryl group conjugation or removal | 0.0% | 0/0 | 0.0% | 0/1 | 0.0% |
| Catalysis of small protein conjugation or removal | 0.0% | 0/0 | 0.0% | 0/2 | 0.0% |
| Other catalysis of small protein conjugation | 0.0% | 0/0 | 0.0% | 0/1 | 0.0% |
| Other catalysis of small protein removal | 0.0% | 0/0 | 0.0% | 0/1 | 0.0% |
| Regulation of translation | 0.0% | 0/0 | 0.0% | 0/7 | 0.0% |

### Supplementary Section 7: Results of relation extraction manual error analysis results for the best model on 20% of documents on the RegulaTome test set

FN: False Negative, FP: False Positive

| PMID | Relationship type | Entity 1 | Entity 2 | Error type | FP/FN |
|---|---|---|---|---|---|
| 1939122 | Complex formation | T13 | T17 | ambiguous keyword | FP |
| 1939122 | Regulation | T5 | T16 | ambiguous keyword | FP |
| 9857197 | Regulation | T15 | T14 | ambiguous keyword | FP |
| 9857197 | Regulation | T28 | T4 | ambiguous keyword | FP |
| 9857197 | Regulation | T5 | T4 | ambiguous keyword | FP |
| 10446169 | Regulation | T23 | T14 | ambiguous keyword | FP |
| 10763828 | Negative regulation | T13 | T15 | ambiguous keyword | FP |
| 10763828 | Regulation of gene expression | T13 | T15 | ambiguous keyword | FP |
| 10763828 | Regulation of gene expression | T16 | T17 | ambiguous keyword | FP |
| 10763828 | Regulation of transcription | T22 | T23 | ambiguous keyword | FP |
| 11099047 | Complex formation | T18 | T20 | ambiguous keyword | FP |
| 11099047 | Complex formation | T19 | T20 | ambiguous keyword | FP |
| 14536086 | Catalysis of methylation | T14 | T12 | ambiguous keyword | FP |
| 14536086 | Catalysis of methylation | T15 | T12 | ambiguous keyword | FP |
| 14981544 | Regulation of transcription | T27 | T28 | ambiguous keyword | FP |
| 14981544 | Regulation of transcription | T3 | T4 | ambiguous keyword | FP |
| 15031289 | Regulation | T10 | T8 | ambiguous keyword | FP |
| 15485920 | Regulation | T24 | T25 | ambiguous keyword | FP |
| 15485920 | Complex formation | T19 | T4 | ambiguous keyword | FP |
| 15485920 | Regulation | T19 | T4 | ambiguous keyword | FP |
| 15886116 | Regulation | T2 | T6 | ambiguous keyword | FP |
| 15886116 | Regulation | T3 | T6 | ambiguous keyword | FP |
| 15886116 | Regulation | T4 | T6 | ambiguous keyword | FP |
| 15886116 | Regulation | T5 | T6 | ambiguous keyword | FP |
| 16282325 | Regulation | T20 | T21 | ambiguous keyword | FP |
| 17572495 | Regulation of transcription | T2 | T3 | ambiguous keyword | FP |
| 17572495 | Regulation of transcription | T1 | T3 | ambiguous keyword | FP |
| 17662961 | Complex formation | T10 | T11 | ambiguous keyword | FP |
| 17662961 | Complex formation | T11 | T12 | ambiguous keyword | FP |
| 17948059 | Catalysis of methylation | T19 | T1 | ambiguous keyword | FP |
| 17948059 | Catalysis of posttranslational modification | T14 | T15 | ambiguous keyword | FP |
| 18094119 | Regulation of transcription | T19 | T18 | ambiguous keyword | FP |
| 18158893 | Positive regulation | T15 | T4 | ambiguous keyword | FP |
| 18158893 | Negative regulation | T6 | T16 | ambiguous keyword | FP |

| PMID | Relationship type | Entity 1 | Entity 2 | Error type | FP/FN |
|---|---|---|---|---|---|
| 18794358 | Regulation | T13 | T14 | ambiguous keyword | FP |
| 18794358 | Regulation | T13 | T15 | ambiguous keyword | FP |
| 20368621 | Regulation of transcription | T9 | T10 | ambiguous keyword | FP |
| 21236256 | Regulation of transcription | T12 | T23 | ambiguous keyword | FP |
| 21236256 | Regulation of transcription | T12 | T24 | ambiguous keyword | FP |
| 21236256 | Regulation of transcription | T11 | T23 | ambiguous keyword | FP |
| 21236256 | Regulation of transcription | T11 | T24 | ambiguous keyword | FP |
| 21266407 | Complex formation | T10 | T11 | ambiguous keyword | FP |
| 21266407 | Regulation | T16 | T4 | ambiguous keyword | FP |
| 21554248 | Regulation of gene expression | T15 | T16 | ambiguous keyword | FP |
| 21554248 | Regulation of gene expression | T14 | T16 | ambiguous keyword | FP |
| 21554248 | Regulation of gene expression | T13 | T16 | ambiguous keyword | FP |
| 21554248 | Regulation of gene expression | T36 | T16 | ambiguous keyword | FP |
| 18995841_22 | Regulation | T361 | T360 | ambiguous keyword | FP |
| 18995841_22 | Regulation | T373 | T374 | ambiguous keyword | FP |
| 18995841_22 | Regulation | T373 | T375 | ambiguous keyword | FP |
| 21347277_5 | Complex formation | T3 | T7 | ambiguous keyword | FP |
| 22086907_3 | Catalysis of ubiquitination | T8 | T13 | ambiguous keyword | FP |
| 22958824_7 | Complex formation | T8 | T9 | ambiguous keyword | FP |
| 23247405_12 | Regulation | T196 | T194 | ambiguous keyword | FP |
| 23247405_12 | Regulation | T196 | T195 | ambiguous keyword | FP |
| 23247405_12 | Regulation | T197 | T194 | ambiguous keyword | FP |
| 23247405_12 | Regulation | T197 | T195 | ambiguous keyword | FP |
| 23615448_14 | Positive regulation | T251 | T248 | ambiguous keyword | FP |
| 23615448_14 | Positive regulation | T250 | T249 | ambiguous keyword | FP |
| 25819761_21 | Complex formation | T341 | T342 | ambiguous keyword | FP |
| 32367036_16 | Regulation | T196 | T195 | ambiguous keyword | FP |
| 9412462_6 | Regulation | T68 | T69 | ambiguous keyword | FP |
| 9412462_6 | Regulation | T68 | T71 | ambiguous keyword | FP |
| 9412462_6 | Regulation | T77 | T78 | ambiguous keyword | FP |
| 9412462_6 | Regulation | T77 | T79 | ambiguous keyword | FP |
| 22086907_3 | Catalysis of posttranslational modification | T22 | T12 | ambiguous keyword | FN |
| 15031289 | Catalysis of posttranslational modification | T10 | T8 | ambiguous keyword | FN |
| 10866691 | Complex formation | T17 | T16 | ambiguous keyword | FN |

| PMID | Relationship type | Entity 1 | Entity 2 | Error type | FP/FN |
|---|---|---|---|---|---|
| 10866691 | Complex formation | T20 | T19 | ambiguous keyword | FN |
| 17108083 | Complex formation | T12 | T13 | ambiguous keyword | FN |
| 18794358 | Complex formation | T13 | T16 | ambiguous keyword | FN |
| 18794358 | Complex formation | T13 | T17 | ambiguous keyword | FN |
| 21236256 | Complex formation | T11 | T12 | ambiguous keyword | FN |
| 23022657_23 | Complex formation | T9 | T8 | ambiguous keyword | FN |
| 23022657_23 | Complex formation | T6 | T7 | ambiguous keyword | FN |
| 9857197 | Complex formation | T15 | T14 | ambiguous keyword | FN |
| 15485920 | Complex formation | T24 | T25 | ambiguous keyword | FN |
| 18794358 | Complex formation | T13 | T14 | ambiguous keyword | FN |
| 18794358 | Complex formation | T13 | T15 | ambiguous keyword | FN |
| 18794358 | Positive regulation | T38 | T40 | ambiguous keyword | FN |
| 18794358 | Positive regulation | T38 | T41 | ambiguous keyword | FN |
| 16282325 | Positive regulation | T20 | T21 | ambiguous keyword | FN |
| 9412462_6 | Positive regulation | T68 | T69 | ambiguous keyword | FN |
| 9412462_6 | Positive regulation | T68 | T71 | ambiguous keyword | FN |
| 9412462_6 | Positive regulation | T77 | T78 | ambiguous keyword | FN |
| 9412462_6 | Positive regulation | T77 | T79 | ambiguous keyword | FN |
| 10763828 | Regulation of gene expression | T22 | T23 | ambiguous keyword | FN |
| 17572495 | Regulation of gene expression | T2 | T3 | ambiguous keyword | FN |
| 17572495 | Regulation of gene expression | T1 | T3 | ambiguous keyword | FN |
| 18158893 | Regulation of gene expression | T17 | T10 | ambiguous keyword | FN |
| 20368621 | Regulation of gene expression | T9 | T10 | ambiguous keyword | FN |
| 18794358 | Regulation of transcription | T38 | T39 | ambiguous keyword | FN |
| 18794358 | Regulation of transcription | T38 | T40 | ambiguous keyword | FN |
| 18794358 | Regulation of transcription | T38 | T41 | ambiguous keyword | FN |
| 21554248 | Regulation of transcription | T15 | T16 | ambiguous keyword | FN |
| 21554248 | Regulation of transcription | T14 | T16 | ambiguous keyword | FN |
| 21554248 | Regulation of transcription | T13 | T16 | ambiguous keyword | FN |
| 21554248 | Regulation of transcription | T36 | T16 | ambiguous keyword | FN |
| 14981544 | Regulation | T33 | T15 | ambiguous keyword | FN |
| 22958824_7 | Regulation | T16 | T15 | ambiguous keyword | FN |
| 1939122 | Negative regulation | T18 | T7 | annotation error | FP |
| 10866691 | Catalysis of phosphorylation | T12 | T13 | annotation error | FP |
| 10866691 | Catalysis of phosphorylation | T30 | T13 | annotation error | FP |
| 10866691 | Regulation | T30 | T14 | annotation error | FP |
| 11983168 | Positive regulation | T5 | T4 | annotation error | FP |
| 12548019 | Positive regulation | T2 | T1 | annotation error | FP |
| 12730668 | Negative regulation | T3 | T5 | annotation error | FP |
| 14536086 | Positive regulation | T23 | T21 | annotation error | FP |

| PMID | Relationship type | Entity 1 | Entity 2 | Error type | FP/FN |
|---|---|---|---|---|---|
| 18794358 | Positive regulation | T25 | T24 | annotation error | FP |
| 18794358 | Regulation | T25 | T23 | annotation error | FP |
| 18794358 | Complex formation | T6 | T10 | annotation error | FP |
| 18794358 | Complex formation | T14 | T16 | annotation error | FP |
| 18794358 | Complex formation | T14 | T17 | annotation error | FP |
| 18794358 | Complex formation | T15 | T16 | annotation error | FP |
| 18794358 | Complex formation | T15 | T17 | annotation error | FP |
| 18794358 | Complex formation | T28 | T32 | annotation error | FP |
| 18794358 | Complex formation | T29 | T31 | annotation error | FP |
| 18794358 | Complex formation | T29 | T32 | annotation error | FP |
| 20457893 | Catalysis of demethylation | T1 | T2 | annotation error | FP |
| 20858899 | Negative regulation | T32 | T33 | annotation error | FP |
| 20858899 | Positive regulation | T32 | T35 | annotation error | FP |
| 21081508 | Negative regulation | T6 | T8 | annotation error | FP |
| 22194618 | Positive regulation | T30 | T10 | annotation error | FP |
| 19637179_9 | Complex formation | T19 | T20 | annotation error | FP |
| 19637179_9 | Complex formation | T4 | T15 | annotation error | FP |
| 22958824_7 | Positive regulation | T5 | T4 | annotation error | FP |
| 21081508 | Catalysis of glycosylation | T6 | T8 | annotation error | FN |
| 17255109 | Complex formation | T20 | T21 | annotation error | FN |
| 18794358 | Complex formation | T18 | T19 | annotation error | FN |
| 20457893 | Complex formation | T12 | T13 | annotation error | FN |
| 29666278 | Complex formation | T10 | T28 | annotation error | FN |
| 18794358 | Complex formation | T25 | T24 | annotation error | FN |
| 18794358 | Complex formation | T23 | T25 | annotation error | FN |
| 15031289 | Negative regulation | T17 | T19 | annotation error | FN |
| 15031289 | Negative regulation | T17 | T20 | annotation error | FN |
| 20858899 | Regulation of degradation | T32 | T33 | annotation error | FN |
| 23240008 | Regulation of transcription | T13 | T12 | annotation error | FN |
| 10770935 | Regulation | T14 | T10 | annotation error | FN |
| 12548019 | Regulation | T2 | T1 | annotation error | FN |
| 20858899 | Regulation | T32 | T35 | annotation error | FN |
| 22958824_7 | Regulation | T17 | T15 | annotation error | FN |
| 14981544 | Negative regulation | T36 | T30 | co-reference resolution | FP |
| 17255109 | Negative regulation | T36 | T35 | co-reference resolution | FP |
| 17255109 | Positive regulation | T20 | T16 | co-reference resolution | FP |
| 17255109 | Positive regulation | T21 | T16 | co-reference resolution | FP |
| 18624796 | Complex formation | T3 | T7 | co-reference resolution | FP |
| 18624796 | Complex formation | T4 | T5 | co-reference resolution | FP |
| 18794358 | Catalysis of acetylation | T29 | T35 | co-reference resolution | FP |
| 18794358 | Catalysis of acetylation | T29 | T36 | co-reference resolution | FP |

| PMID | Relationship type | Entity 1 | Entity 2 | Error type | FP/FN |
|---|---|---|---|---|---|
| 18794358 | Complex formation | T26 | T27 | co-reference resolution | FP |
| 19503082 | Regulation | T18 | T19 | co-reference resolution | FP |
| 20129058 | Complex formation | T18 | T15 | co-reference resolution | FP |
| 20406818 | Catalysis of ubiquitination | T6 | T8 | co-reference resolution | FP |
| 20406818 | Complex formation | T25 | T27 | co-reference resolution | FP |
| 20624928 | Positive regulation | T20 | T3 | co-reference resolution | FP |
| 21236256 | Regulation of transcription | T6 | T18 | co-reference resolution | FP |
| 21236256 | Regulation of transcription | T6 | T19 | co-reference resolution | FP |
| 21236256 | Regulation of transcription | T5 | T18 | co-reference resolution | FP |
| 21236256 | Regulation of transcription | T5 | T19 | co-reference resolution | FP |
| 21266407 | Regulation | T11 | T23 | co-reference resolution | FP |
| 27578797 | Regulation | T1 | T12 | co-reference resolution | FP |
| 29666278 | Negative regulation | T3 | T13 | co-reference resolution | FP |
| 29666278 | Regulation of degradation | T3 | T13 | co-reference resolution | FP |
| 11238463_35 | Negative regulation | T272 | T269 | co-reference resolution | FP |
| 11238463_35 | Negative regulation | T272 | T270 | co-reference resolution | FP |
| 11238463_35 | Negative regulation | T273 | T270 | co-reference resolution | FP |
| 18995841_22 | Negative regulation | T371 | T370 | co-reference resolution | FP |
| 19684574_7 | Regulation | T7 | T19 | co-reference resolution | FP |
| 23022657_23 | Complex formation | T6 | T10 | co-reference resolution | FP |
| 26984517_50 | Negative regulation | T311 | T314 | co-reference resolution | FP |
| 26984517_50 | Negative regulation | T311 | T315 | co-reference resolution | FP |
| 26984517_50 | Negative regulation | T312 | T314 | co-reference resolution | FP |
| 17615152 | Catalysis of phosphorylation | T42 | T11 | co-reference resolution | FN |
| 10446169 | Complex formation | T12 | T13 | co-reference resolution | FN |
| 16282325 | Complex formation | T17 | T18 | co-reference resolution | FN |
| 16282325 | Complex formation | T17 | T19 | co-reference resolution | FN |
| 17255109 | Complex formation | T37 | T38 | co-reference resolution | FN |
| 18794358 | Complex formation | T14 | T15 | co-reference resolution | FN |
| 18794358 | Complex formation | T16 | T17 | co-reference resolution | FN |
| 18794358 | Complex formation | T28 | T29 | co-reference resolution | FN |
| 17615152 | Negative regulation | T12 | T1 | co-reference resolution | FN |
| 17615152 | Negative regulation | T13 | T1 | co-reference resolution | FN |
| 26984517_50 | Negative regulation | T310 | T314 | co-reference resolution | FN |
| 26984517_50 | Negative regulation | T310 | T315 | co-reference resolution | FN |
| 1939122 | Positive regulation | T29 | T15 | co-reference resolution | FN |
| 16282325 | Positive regulation | T25 | T28 | co-reference resolution | FN |

| PMID | Relationship type | Entity 1 | Entity 2 | Error type | FP/FN |
|---|---|---|---|---|---|
| 21554248 | Positive regulation | T21 | T19 | co-reference resolution | FN |
| 21554248 | Positive regulation | T35 | T42 | co-reference resolution | FN |
| 21554248 | Positive regulation | T20 | T19 | co-reference resolution | FN |
| 21554248 | Positive regulation | T8 | T42 | co-reference resolution | FN |
| 17572495 | Positive regulation | T30 | T29 | co-reference resolution | FN |
| 17572495 | Positive regulation | T31 | T29 | co-reference resolution | FN |
| 17572495 | Positive regulation | T32 | T29 | co-reference resolution | FN |
| 17615152 | Regulation of gene expression | T12 | T1 | co-reference resolution | FN |
| 17615152 | Regulation of gene expression | T13 | T1 | co-reference resolution | FN |
| 11983168 | Regulation | T19 | T22 | co-reference resolution | FN |
| 11983168 | Regulation | T19 | T23 | co-reference resolution | FN |
| 14752096 | Regulation | T6 | T18 | co-reference resolution | FN |
| 18094119 | Regulation | T9 | T6 | co-reference resolution | FN |
| 18094119 | Regulation | T9 | T7 | co-reference resolution | FN |
| 27578797 | Regulation | T1 | T2 | co-reference resolution | FN |
| 1939122 | Positive regulation | T25 | T7 | convoluted text excerpt | FP |
| 1939122 | Positive regulation | T26 | T7 | convoluted text excerpt | FP |
| 1939122 | Positive regulation | T33 | T23 | convoluted text excerpt | FP |
| 1939122 | Positive regulation | T33 | T24 | convoluted text excerpt | FP |
| 1939122 | Regulation | T30 | T15 | convoluted text excerpt | FP |
| 2981876 | Complex formation | T14 | T17 | convoluted text excerpt | FP |
| 2981876 | Complex formation | T15 | T17 | convoluted text excerpt | FP |
| 2981876 | Complex formation | T16 | T17 | convoluted text excerpt | FP |
| 2981876 | Negative regulation | T23 | T8 | convoluted text excerpt | FP |
| 2981876 | Negative regulation | T23 | T9 | convoluted text excerpt | FP |
| 10763828 | Negative regulation | T26 | T25 | convoluted text excerpt | FP |
| 10763828 | Regulation of gene expression | T26 | T25 | convoluted text excerpt | FP |
| 10770935 | Positive regulation | T18 | T7 | convoluted text excerpt | FP |
| 10770935 | Regulation of transcription | T18 | T7 | convoluted text excerpt | FP |
| 11099047 | Catalysis of deacetylation | T7 | T8 | convoluted text excerpt | FP |
| 11598127 | Catalysis of ubiquitination | T6 | T7 | convoluted text excerpt | FP |
| 11598127 | Positive regulation | T18 | T22 | convoluted text excerpt | FP |
| 11687605 | Negative regulation | T2 | T5 | convoluted text excerpt | FP |
| 11687605 | Regulation of transcription | T2 | T5 | convoluted text excerpt | FP |
| 11983168 | Regulation | T20 | T22 | convoluted text excerpt | FP |
| 11983168 | Regulation | T20 | T23 | convoluted text excerpt | FP |
| 12730668 | Negative regulation | T19 | T18 | convoluted text excerpt | FP |
| 12730668 | Positive regulation | T20 | T18 | convoluted text excerpt | FP |
| 12730668 | Positive regulation | T25 | T26 | convoluted text excerpt | FP |
| 12730668 | Regulation of transcription | T19 | T18 | convoluted text excerpt | FP |
| 14752096 | Catalysis of acetylation | T5 | T16 | convoluted text excerpt | FP |

| PMID | Relationship type | Entity 1 | Entity 2 | Error type | FP/FN |
|---|---|---|---|---|---|
| 15031289 | Negative regulation | T22 | T23 | convoluted text excerpt | FP |
| 15031289 | Negative regulation | T22 | T24 | convoluted text excerpt | FP |
| 16282325 | Regulation | T29 | T30 | convoluted text excerpt | FP |
| 17108083 | Catalysis of ubiquitination | T14 | T18 | convoluted text excerpt | FP |
| 17108083 | Catalysis of ubiquitination | T27 | T31 | convoluted text excerpt | FP |
| 17255109 | Complex formation | T18 | T19 | convoluted text excerpt | FP |
| 17572495 | Positive regulation | T6 | T7 | convoluted text excerpt | FP |
| 18794358 | Complex formation | T6 | T11 | convoluted text excerpt | FP |
| 18794358 | Complex formation | T6 | T12 | convoluted text excerpt | FP |
| 18794358 | Complex formation | T8 | T10 | convoluted text excerpt | FP |
| 18794358 | Complex formation | T8 | T11 | convoluted text excerpt | FP |
| 18794358 | Complex formation | T8 | T12 | convoluted text excerpt | FP |
| 18794358 | Complex formation | T10 | T11 | convoluted text excerpt | FP |
| 19503082 | Positive regulation | T23 | T6 | convoluted text excerpt | FP |
| 20368621 | Negative regulation | T9 | T18 | convoluted text excerpt | FP |
| 20512148 | Positive regulation | T20 | T2 | convoluted text excerpt | FP |
| 20512148 | Regulation of gene expression | T20 | T2 | convoluted text excerpt | FP |
| 21236256 | Positive regulation | T15 | T17 | convoluted text excerpt | FP |
| 21554248 | Regulation of transcription | T11 | T42 | convoluted text excerpt | FP |
| 21554248 | Positive regulation | T23 | T24 | convoluted text excerpt | FP |
| 21554248 | Positive regulation | T27 | T38 | convoluted text excerpt | FP |
| 22908299 | Regulation | T20 | T19 | convoluted text excerpt | FP |
| 27578797 | Negative regulation | T10 | T16 | convoluted text excerpt | FP |
| 29666278 | Catalysis of ubiquitination | T20 | T19 | convoluted text excerpt | FP |
| 29666278 | Catalysis of ubiquitination | T21 | T19 | convoluted text excerpt | FP |
| 29666278 | Catalysis of ubiquitination | T22 | T19 | convoluted text excerpt | FP |
| 29666278 | Regulation of degradation | T20 | T19 | convoluted text excerpt | FP |
| 29666278 | Regulation of degradation | T21 | T19 | convoluted text excerpt | FP |
| 29666278 | Regulation of degradation | T22 | T19 | convoluted text excerpt | FP |
| 18995841_22 | Positive regulation | T354 | T357 | convoluted text excerpt | FP |
| 18995841_22 | Positive regulation | T356 | T357 | convoluted text excerpt | FP |
| 19637179_9 | Regulation | T19 | T10 | convoluted text excerpt | FP |
| 21951725_19 | Complex formation | T230 | T237 | convoluted text excerpt | FP |
| 22086907_3 | Negative regulation | T20 | T30 | convoluted text excerpt | FP |
| 26984517_50 | Negative regulation | T302 | T301 | convoluted text excerpt | FP |
| 17255109 | Catalysis of phosphoryl group conjugation or removal | T1 | T2 | convoluted text excerpt | FN |
| 19503082 | Catalysis of phosphorylation | T24 | T9 | convoluted text excerpt | FN |
| 23240008 | Catalysis of phosphorylation | T9 | T10 | convoluted text excerpt | FN |

| PMID | Relationship type | Entity 1 | Entity 2 | Error type | FP/FN |
|---|---|---|---|---|---|
| 17018294 | Catalysis of SUMOylation | T11 | T9 | convoluted text excerpt | FN |
| 17255109 | Catalysis of ubiquitination | T34 | T35 | convoluted text excerpt | FN |
| 12730668 | Complex formation | T23 | T24 | convoluted text excerpt | FN |
| 21554248 | Complex formation | T11 | T9 | convoluted text excerpt | FN |
| 21554248 | Complex formation | T9 | T35 | convoluted text excerpt | FN |
| 21554248 | Complex formation | T21 | T22 | convoluted text excerpt | FN |
| 21554248 | Complex formation | T20 | T22 | convoluted text excerpt | FN |
| 32367036_16 | Complex formation | T193 | T194 | convoluted text excerpt | FN |
| 11598127 | Negative regulation | T14 | T16 | convoluted text excerpt | FN |
| 11598127 | Negative regulation | T13 | T16 | convoluted text excerpt | FN |
| 12730668 | Negative regulation | T30 | T16 | convoluted text excerpt | FN |
| 12730668 | Negative regulation | T17 | T16 | convoluted text excerpt | FN |
| 18158893 | Negative regulation | T8 | T18 | convoluted text excerpt | FN |
| 19503082 | Negative regulation | T11 | T13 | convoluted text excerpt | FN |
| 21236256 | Negative regulation | T10 | T23 | convoluted text excerpt | FN |
| 21236256 | Negative regulation | T10 | T24 | convoluted text excerpt | FN |
| 26984517_50 | Negative regulation | T303 | T302 | convoluted text excerpt | FN |
| 32367036_16 | Negative regulation | T193 | T194 | convoluted text excerpt | FN |
| 20512148 | Negative regulation | T24 | T4 | convoluted text excerpt | FN |
| 1939122 | Positive regulation | T22 | T15 | convoluted text excerpt | FN |
| 12548019 | Positive regulation | T14 | T17 | convoluted text excerpt | FN |
| 15031289 | Positive regulation | T22 | T24 | convoluted text excerpt | FN |
| 27578797 | Positive regulation | T10 | T16 | convoluted text excerpt | FN |
| 1939122 | Positive regulation | T30 | T15 | convoluted text excerpt | FN |
| 29666278 | Regulation of degradation | T6 | T19 | convoluted text excerpt | FN |
| 18158893 | Regulation of gene expression | T17 | T11 | convoluted text excerpt | FN |
| 19503082 | Regulation of gene expression | T11 | T13 | convoluted text excerpt | FN |
| 26984517_50 | Regulation of gene expression | T303 | T302 | convoluted text excerpt | FN |
| 20406818 | Regulation of transcription | T8 | T12 | convoluted text excerpt | FN |
| 7524088 | Regulation | T4 | T7 | convoluted text excerpt | FN |
| 9857197 | Regulation | T10 | T8 | convoluted text excerpt | FN |
| 9857197 | Regulation | T10 | T9 | convoluted text excerpt | FN |
| 15886116 | Regulation | T28 | T13 | convoluted text excerpt | FN |
| 17108083 | Regulation | T14 | T16 | convoluted text excerpt | FN |
| 17108083 | Regulation | T14 | T17 | convoluted text excerpt | FN |
| 18094119 | Regulation | T17 | T14 | convoluted text excerpt | FN |
| 18158893 | Regulation | T2 | T13 | convoluted text excerpt | FN |
| 18158893 | Regulation | T18 | T9 | convoluted text excerpt | FN |

| PMID | Relationship type | Entity 1 | Entity 2 | Error type | FP/FN |
|---|---|---|---|---|---|
| 20406818 | Regulation | T1 | T2 | convoluted text excerpt | FN |
| 21236256 | Regulation | T1 | T16 | convoluted text excerpt | FN |
| 21236256 | Regulation | T1 | T17 | convoluted text excerpt | FN |
| 21236256 | Regulation | T13 | T14 | convoluted text excerpt | FN |
| 23240008 | Regulation | T20 | T19 | convoluted text excerpt | FN |
| 29666278 | Regulation | T8 | T25 | convoluted text excerpt | FN |
| 18995841_22 | Regulation | T364 | T367 | convoluted text excerpt | FN |
| 18995841_22 | Regulation | T365 | T367 | convoluted text excerpt | FN |
| 18995841_22 | Regulation | T366 | T367 | convoluted text excerpt | FN |
| 25819761_21 | Regulation | T346 | T345 | convoluted text excerpt | FN |
| 18158893 | Regulation | T6 | T16 | convoluted text excerpt | FN |
| 26984517_50 | Regulation | T302 | T301 | convoluted text excerpt | FN |
| 21554248 | Regulation | T23 | T24 | convoluted text excerpt | FN |
| 21554248 | Regulation | T27 | T38 | convoluted text excerpt | FN |
| 11099047 | Catalysis of deacetylation | T12 | T13 | model error | FP |
| 11099047 | Catalysis of deacetylation | T12 | T14 | model error | FP |
| 11099047 | Complex formation | T8 | T11 | model error | FP |
| 11099047 | Catalysis of deacetylation | T20 | T21 | model error | FP |
| 17662961 | Complex formation | T1 | T23 | model error | FP |
| 18094119 | Negative regulation | T9 | T7 | model error | FP |
| 18158893 | Regulation of transcription | T17 | T10 | model error | FP |
| 20406818 | Positive regulation | T16 | T17 | model error | FP |
| 20512148 | Regulation | T24 | T4 | model error | FP |
| 20858899 | Catalysis of small protein conjugation | T36 | T37 | model error | FP |
| 22194618 | Complex formation | T28 | T6 | model error | FP |
| 21685908_17 | Regulation | T11 | T10 | model error | FP |
| 21951725_19 | Complex formation | T244 | T245 | model error | FP |
| 23022657_23 | Complex formation | T1 | T2 | model error | FP |
| 23247405_9 | Complex formation | T101 | T1 | model error | FP |
| 23247405_9 | Complex formation | T101 | T104 | model error | FP |
| 25916855_48 | Positive regulation | T161 | T160 | model error | FP |
| 18794358 | Catalysis of acetylation | T1 | T3 | model error | FN |
| 18794358 | Catalysis of acetylation | T2 | T3 | model error | FN |
| 12730668 | Catalysis of deacetylation | T5 | T4 | model error | FN |
| 11341840 | Catalysis of methylation | T7 | T6 | model error | FN |

| PMID | Relationship type | Entity 1 | Entity 2 | Error type | FP/FN |
|---|---|---|---|---|---|
| 11341840 | Catalysis of methylation | T17 | T16 | model error | FN |
| 15145825 | Catalysis of methylation | T7 | T6 | model error | FN |
| 15145825 | Catalysis of methylation | T15 | T18 | model error | FN |
| 15145825 | Catalysis of methylation | T14 | T13 | model error | FN |
| 17018294 | Catalysis of SUMOylation | T5 | T6 | model error | FN |
| 10770935 | Complex formation | T3 | T7 | model error | FN |
| 11099047 | Complex formation | T8 | T9 | model error | FN |
| 18794358 | Complex formation | T19 | T21 | model error | FN |
| 18794358 | Complex formation | T19 | T22 | model error | FN |
| 19637179_9 | Complex formation | T14 | T29 | model error | FN |
| 10763828 | Negative regulation | T33 | T21 | model error | FN |
| 22194618 | Negative regulation | T5 | T28 | model error | FN |
| 22194618 | Negative regulation | T5 | T7 | model error | FN |
| 19637179_9 | Negative regulation | T19 | T10 | model error | FN |
| 10763828 | Negative regulation | T16 | T17 | model error | FN |
| 12548019 | Positive regulation | T11 | T8 | model error | FN |
| 23240008 | Regulation of transcription | T3 | T4 | model error | FN |
| 11099047 | Regulation | T20 | T21 | model error | FN |
| 9857197 | Regulation | T1 | T26 | model error | FN |
| 14752096 | Regulation | T1 | T7 | model error | FN |
| 18094119 | Regulation | T26 | T22 | model error | FN |
| 20858899 | Regulation | T32 | T34 | model error | FN |
| 22958824_7 | Regulation | T5 | T4 | model error | FN |
| 25819761_21 | Regulation | T340 | T342 | model error | FN |
| 18094119 | Regulation | T24 | T22 | model error | FN |
| 20406818 | Regulation | T16 | T17 | model error | FN |
| 25916855_48 | Regulation | T161 | T160 | model error | FN |
| 18094119 | Regulation | T25 | T22 | model error | FN |
| 15031289 | Catalysis of phosphorylation | T15 | T16 | rare keyword | FN |
| 2002555 | Complex formation | T5 | T4 | rare keyword | FN |
| 14981544 | Complex formation | T27 | T28 | rare keyword | FN |
| 18624796 | Complex formation | T20 | T21 | rare keyword | FN |
| 18624796 | Complex formation | T19 | T21 | rare keyword | FN |
| 20129058 | Complex formation | T14 | T15 | rare keyword | FN |
| 22086907_3 | Complex formation | T5 | T4 | rare keyword | FN |
| 22219378_47 | Complex formation | T704 | T706 | rare keyword | FN |
| 26984517_50 | Complex formation | T304 | T305 | rare keyword | FN |
| 26984517_50 | Complex formation | T306 | T305 | rare keyword | FN |

| PMID | Relationship type | Entity 1 | Entity 2 | Error type | FP/FN |
|---|---|---|---|---|---|
| 26984517_50 | Complex formation | T307 | T309 | rare keyword | FN |
| 26984517_50 | Complex formation | T311 | T312 | rare keyword | FN |
| 26984517_50 | Complex formation | T311 | T313 | rare keyword | FN |
| 18094119 | Negative regulation | T13 | T14 | rare keyword | FN |
| 18094119 | Negative regulation | T24 | T23 | rare keyword | FN |
| 18094119 | Negative regulation | T25 | T23 | rare keyword | FN |
| 18094119 | Negative regulation | T26 | T23 | rare keyword | FN |
| 22086907_3 | Negative regulation | T9 | T14 | rare keyword | FN |
| 12548019 | Positive regulation | T23 | T11 | rare keyword | FN |
| 14536086 | Positive regulation | T1 | T2 | rare keyword | FN |
| 14536086 | Positive regulation | T27 | T29 | rare keyword | FN |
| 14536086 | Positive regulation | T26 | T29 | rare keyword | FN |
| 17615152 | Positive regulation | T19 | T7 | rare keyword | FN |
| 17615152 | Positive regulation | T22 | T7 | rare keyword | FN |
| 21236256 | Positive regulation | T8 | T20 | rare keyword | FN |
| 21236256 | Positive regulation | T8 | T21 | rare keyword | FN |
| 27578797 | Positive regulation | T6 | T7 | rare keyword | FN |
| 29666278 | Positive regulation | T6 | T19 | rare keyword | FN |
| 23615448_14 | Positive regulation | T245 | T246 | rare keyword | FN |
| 22086907_3 | Regulation of degradation | T9 | T14 | rare keyword | FN |
| 17615152 | Regulation of gene expression | T19 | T7 | rare keyword | FN |
| 17615152 | Regulation of gene expression | T22 | T7 | rare keyword | FN |
| 17572495 | Regulation of transcription | T33 | T35 | rare keyword | FN |
| 17572495 | Regulation of transcription | T34 | T35 | rare keyword | FN |
| 21554248 | Regulation of transcription | T1 | T45 | rare keyword | FN |
| 21554248 | Regulation of transcription | T35 | T42 | rare keyword | FN |
| 21554248 | Regulation of transcription | T8 | T42 | rare keyword | FN |
| 7524088 | Regulation | T14 | T10 | rare keyword | FN |
| 7524088 | Regulation | T14 | T11 | rare keyword | FN |
| 7524088 | Regulation | T14 | T12 | rare keyword | FN |
| 10446169 | Regulation | T5 | T6 | rare keyword | FN |
| 10446169 | Regulation | T4 | T6 | rare keyword | FN |
| 17018294 | Regulation | T3 | T1 | rare keyword | FN |
| 18094119 | Regulation | T19 | T18 | rare keyword | FN |
| 20129058 | Regulation | T3 | T4 | rare keyword | FN |
| 20129058 | Regulation | T17 | T12 | rare keyword | FN |
| 20129058 | Regulation | T20 | T2 | rare keyword | FN |
| 20368621 | Regulation | T3 | T4 | rare keyword | FN |

| PMID | Relationship type | Entity 1 | Entity 2 | Error type | FP/FN |
|---|---|---|---|---|---|
| 20368621 | Regulation | T5 | T4 | rare keyword | FN |
| 20512148 | Regulation | T28 | T27 | rare keyword | FN |
| 27578797 | Regulation | T4 | T22 | rare keyword | FN |
| 27578797 | Regulation | T5 | T23 | rare keyword | FN |
| 27578797 | Regulation | T19 | T24 | rare keyword | FN |
| 27578797 | Regulation | T8 | T19 | rare keyword | FN |
| 23615448_14 | Regulation | T253 | T254 | rare keyword | FN |
| 23615448_14 | Regulation | T261 | T262 | rare keyword | FN |
| 9008702_22 | Regulation | T104 | T108 | rare keyword | FN |

### Supplementary Section 8: Error confusion for False Positives in the RegulaTome test set

| Confusion category | #FP |
|---|---|
| ***Negative predicted as Complex formation*** | ***244*** |
| ***Negative predicted as Regulation*** | ***156*** |
| ***Negative predicted as Positive Regulation*** | ***130*** |
| ***Negative predicted as Negative Regulation*** | ***113*** |
| ***Negative predicted as Regulation of transcription*** | ***44*** |
| ***Negative predicted as Catalysis of ubiquitination*** | ***34*** |
| ***Negative predicted as Regulation of degradation*** | ***33*** |
| Regulation predicted as Positive Regulation | 26 |
| Positive regulation predicted as Regulation | 19 |
| ***Negative predicted as Catalysis of phosphorylation*** | ***15*** |
| Regulation of transcription predicted as Regulation of gene expression | 13 |
| ***Negative predicted as Catalysis of methylation*** | ***14*** |
| ***Negative predicted as Regulation of gene expression*** | ***16*** |
| ***Negative predicted as Catalysis of dephosphorylation*** | ***12*** |
| Regulation predicted as Negative Regulation | 12 |
| ***Negative predicted as Catalysis of acetylation*** | ***11*** |
| Complex formation predicted as Regulation | 10 |
| Regulation of gene expression predicted as Regulation of transcription | 10 |
| Negative regulation predicted as Regulation | 9 |
| ***Negative predicted as Catalysis of posttranslational modification*** | ***7*** |
| Negative regulation predicted as Positive Regulation | 7 |
| Positive regulation predicted as Negative Regulation | 7 |
| Catalysis of posttranslational modification predicted as Complex formation | 6 |
| Regulation predicted as Regulation of transcription | 6 |
| Catalysis of posttranslational modification predicted as Catalysis of phosphorylation | 5 |
| Catalysis of posttranslational modification predicted as Catalysis of ubiquitination | 5 |
| Regulation of translation predicted as Regulation of transcription | 5 |
| Regulation predicted as Complex formation | 5 |
| Catalysis of other small molecule conjugation or removal predicted as Catalysis of demethylation | 4 |
| Complex formation predicted as Negative regulation | 4 |
| Complex Formation predicted as Positive Regulation | 4 |
| ***Negative predicted as Catalysis of deacetylation*** | ***4*** |
| ***Negative predicted as Catalysis of palmitoylation*** | ***4*** |
| Negative regulation predicted as Complex formation | 4 |

| Confusion category | #FP |
|---|---|
| Catalysis of phosphorylation predicted as Catalysis of dephosphorylation | 3 |
| Catalysis of small protein conjugation predicted as Catalysis of SUMOylation | 3 |
| ***Negative predicted as Catalysis of demethylation*** | ***3*** |
| Regulation of degradation predicted as Negative Regulation | 3 |
| Regulation predicted as Catalysis of posttranslational modification | 3 |
| Regulation predicted as Regulation of gene expression | 1 |
| Regulation predicted as Regulation (reverse) | 3 |
| Negative Regulation predicted as Negative Regulation (reverse) | 2 |
| Catalysis of acetylation predicted as Positive regulation | 2 |
| ***Negative predicted as Catalysis of ADP-ribosylation*** | ***2*** |
| ***Negative predicted as Catalysis of glycosylation*** | ***2*** |
| ***Negative predicted as Catalysis of SUMOylation*** | ***2*** |
| Other catalysis of small molecule conjugation predicted as Catalysis of glycosylation | 2 |
| Catalysis of acylation predicted as Catalysis of palmitoylation | 1 |
| Catalysis of demethylation predicted as Catalysis of deacetylation | 1 |
| Catalysis of demethylation predicted as Positive regulation | 1 |
| Catalysis of glycosylation predicted as Negative Regulation | 1 |
| Catalysis of methylation predicted as Negative regulation | 1 |
| Catalysis of neddylation predicted as Complex formation | 1 |
| Catalysis of posttranslational modification predicted as Regulation | 1 |
| Catalysis of ubiquitination predicted as Catalysis of small protein conjugation | 1 |
| Negative regulation predicted as Regulation of gene expression | 1 |
| Other catalysis of small protein removal predicted as Catalysis of deubiquitination | 1 |
| Positive regulation predicted as Complex formation | 1 |
| Regulation of degradation predicted as Complex formation | 1 |
| Regulation of transcription predicted as Catalysis of methylation | 1 |
| Regulation of transcription predicted as Positive Regulation | 1 |
| Regulation predicted as Catalysis of deacetylation | 1 |
| Regulation predicted as Other catalysis of small molecule conjugation | 1 |
| Regulation predicted as Regulation of degradation | 1 |
| Regulation of translation predicted as Regulation of gene expression | 2 |
| **Total** | **1048** |

### Supplementary Section 9: Error confusion for False Negatives in the RegulaTome test set

| Confusion category | #FN |
|---|---|
| ***Complex formation predicted as Negative*** | ***237*** |
| ***Regulation predicted as Negative*** | ***201*** |
| ***Positive Regulation predicted as Negative*** | ***137*** |
| ***Negative regulation predicted as Negative*** | ***128*** |
| ***Regulation of transcription predicted as Negative*** | ***52*** |
| ***Regulation of gene expression predicted as Negative*** | ***38*** |
| ***Catalysis of phosphorylation predicted as Negative*** | ***30*** |
| Regulation predicted as Positive Regulation | 26 |
| ***Catalysis of posttranslational modification predicted as Negative*** | ***21*** |
| Positive regulation predicted as Regulation | 19 |
| ***Catalysis of ubiquitination predicted as Negative*** | ***17*** |
| ***Catalysis of methylation predicted as Negative*** | ***15*** |
| Regulation of transcription predicted as Regulation of gene expression | 14 |
| ***Catalysis of dephosphorylation predicted as Negative*** | ***13*** |
| Regulation predicted as Negative Regulation | 12 |
| ***Catalysis of acetylation predicted as Negative*** | ***10*** |
| Complex formation predicted as Regulation | 9 |
| Negative regulation predicted as Regulation | 9 |
| ***Regulation of degradation predicted as Negative*** | ***9*** |
| Regulation of gene expression predicted as Regulation of transcription | 10 |
| ***Catalysis of SUMOylation predicted as Negative*** | ***7*** |
| Negative regulation predicted as Positive regulation | 7 |
| Positive regulation predicted as Negative Regulation | 7 |
| Regulation predicted as Complex formation | 7 |
| ***Catalysis of deneddylation predicted as Negative*** | ***6*** |
| ***Catalysis of glycosylation predicted as Negative*** | ***6*** |
| Catalysis of posttranslational modification predicted as Complex formation | 6 |
| Regulation predicted as Regulation of transcription | 6 |
| ***Catalysis of palmitoylation predicted as Negative*** | ***5*** |
| Catalysis of posttranslational modification predicted as Catalysis of phosphorylation | 5 |
| Catalysis of posttranslational modification predicted as Catalysis of ubiquitination | 5 |
| Complex formation predicted as Positive Regulation | 5 |
| Negative regulation predicted as Complex Formation | 5 |
| Regulation of translation predicted as Regulation of transcription | 5 |
| ***Catalysis of ADP-ribosylation predicted as Negative*** | ***4*** |
| ***Catalysis of deacetylation predicted as Negative*** | ***4*** |

| Confusion category | #FN |
|---|---|
| Catalysis of other small molecule conjugation or removal predicted as Catalysis of demethylation | 4 |
| Complex formation predicted as Negative regulation | 4 |
| ***Catalysis of deubiquitination predicted as Negative*** | ***3*** |
| Catalysis of phosphorylation predicted as Catalysis of dephosphorylation | 3 |
| Catalysis of small protein conjugation predicted as Catalysis of SUMOylation | 3 |
| Regulation of degradation predicted as Negative Regulation | 3 |
| Regulation predicted as Catalysis of posttranslational modification | 3 |
| Regulation predicted as Regulation (reverse) | 3 |
| Catalysis of acetylation predicted as Positive regulation | 2 |
| ***Catalysis of other small molecule conjugation or removal predicted as Negative*** | ***2*** |
| ***Catalysis of small protein conjugation or removal predicted as Negative*** | ***2*** |
| Negative Regulation predicted as Negative Regulation (reverse) | 2 |
| Negative regulation predicted as Regulation of gene expression | 1 |
| Other catalysis of small molecule conjugation predicted as Catalysis of glycosylation | 2 |
| Regulation of gene expression predicted as Complex formation | 2 |
| Regulation of translation predicted as Regulation of gene expression | 2 |
| Catalysis of acylation predicted as Catalysis of palmitoylation | 1 |
| Catalysis of demethylation predicted as Catalysis of deacetylation | 1 |
| ***Catalysis of demethylation predicted as Negative*** | ***1*** |
| Catalysis of glycosylation predicted as Negative Regulation | 1 |
| Catalysis of methylation predicted as Negative regulation | 1 |
| Catalysis of neddylation predicted as Complex formation | 1 |
| ***Catalysis of phosphoryl group conjugation or removal predicted as Negative*** | ***1*** |
| Catalysis of posttranslational modification predicted as Regulation | 1 |
| ***Catalysis of small protein conjugation predicted as Negative*** | ***1*** |
| Complex formation predicted as Regulation of transcription | 1 |
| Negative regulation predicted as Regulation of transcription | 1 |
| Other catalysis of small molecule conjugation predicted as Catalysis of posttranslational modification | 1 |
| ***Other catalysis of small protein conjugation predicted as Negative*** | ***1*** |
| Other catalysis of small protein removal predicted as Catalysis of deubiquitination | 1 |
| Positive regulation predicted as Catalysis of palmitoylation | 1 |
| Positive regulation predicted as Complex formation | 1 |
| Regulation of degradation predicted as Complex formation | 1 |
| Regulation of transcription predicted as Catalysis of methylation | 1 |
| Regulation of transcription predicted as Positive regulation | 1 |
| Regulation predicted as Catalysis of deacetylation | 1 |
| Regulation predicted as Other catalysis of small molecule conjugation | 1 |
| Regulation predicted as Regulation of gene expression | 1 |
| **Total** | **1,160** |

### Supplementary Section 10: Regulatory network evaluation

#### *Correctly assigned directed interactions*

$TP_{\text{Catalysis of deubiquitination}}$ + $TP_{\text{Catalysis of demethylation}}$ + $TP_{\text{Catalysis of ubiquitination}}$ + $TP_{\text{Catalysis of phosphorylation}}$ + $TP_{\text{Catalysis of dephosphorylation}}$ + $TP_{\text{Catalysis of methylation}}$ + $TP_{\text{Catalysis of neddylation}}$ + $TP_{\text{Negative regulation}}$ + $TP_{\text{Positive regulation}}$ + $TP_{\text{Regulation of degradation}}$ + $TP_{\text{Regulation of transcription}}$ + $TP_{\text{Catalysis of acetylation}}$ + $TP_{\text{Catalysis of palmitoylation}}$ + $TP_{\text{Regulation of gene expression}}$ + $TP_{\text{Catalysis of SUMOylation}}$ + $TP_{\text{Catalysis of deacetylation}}$ + $TP_{\text{Regulation}}$ + $TP_{\text{Catalysis of ADP-ribosylation}}$ + $TP_{\text{Other catalysis of small molecule conjugation}}$ + $TP_{\text{Catalysis of small protein conjugation}}$ + $TP_{\text{Catalysis of posttranslational modification}}$ + $TP_{\text{Catalysis of glycosylation}}$ + Regulation predicted as Positive Regulation + Positive regulation predicted as Regulation + Regulation of transcription predicted as Regulation of gene expression + Regulation predicted as Negative Regulation + Regulation of gene expression predicted as Regulation of transcription + Negative regulation predicted as Regulation + Negative regulation predicted as Positive Regulation + Positive regulation predicted as Negative Regulation + Regulation predicted as Regulation of transcription + Catalysis of posttranslational modification predicted as Catalysis of phosphorylation + Catalysis of posttranslational modification predicted as Catalysis of ubiquitination + Regulation of translation predicted as Regulation of transcription + Catalysis of other small molecule conjugation or removal predicted as Catalysis of demethylation + Catalysis of phosphorylation predicted as Catalysis of dephosphorylation + Catalysis of small protein conjugation predicted as Catalysis of SUMOylation + Regulation of degradation predicted as Negative Regulation + Regulation predicted as Catalysis of posttranslational modification + Regulation predicted as Regulation of gene expression + Catalysis of acetylation predicted as Positive regulation + Other catalysis of small molecule conjugation predicted as Catalysis of glycosylation + Catalysis of acylation predicted as Catalysis of palmitoylation + Catalysis of demethylation predicted as Catalysis of deacetylation + Catalysis of demethylation predicted as Positive regulation + Catalysis of glycosylation predicted as Negative Regulation + Catalysis of methylation predicted as Negative regulation + Catalysis of posttranslational modification predicted as Regulation + Catalysis of ubiquitination predicted as Catalysis of small protein conjugation + Negative regulation predicted as Regulation of gene expression + Other catalysis of small protein removal predicted as Catalysis of deubiquitination + Regulation of transcription predicted as Catalysis of methylation + Regulation of transcription predicted as Positive Regulation + Regulation predicted as Catalysis of deacetylation + Regulation predicted as Other catalysis of small molecule conjugation + Regulation predicted as Regulation of degradation + Regulation of translation predicted as Regulation of gene expression = 12 + 16 + 77 + 65 + 32 + 35 + 1 + 263 + 276 + 36 + 102 + 17 + 7 + 48 + 7 + 5 + 223 + 2 + 1 + 1 + 6 + 1 + 26 + 19 + 13 + 12 + 10 + 9 + 7 + 7 + 6 + 5 + 5 + 5 + 4 + 3 + 3 + 3 + 3 + 1 + 2 + 2 + 1 + 1 + 1 + 1 + 1 + 1 + 1 + 1 + 1 + 1 + 1 + 1 + 1 + 1 + 2 = 1394

#### *Failed to assign a directed interaction, where there should be one*

Regulation predicted as Negative + Positive Regulation predicted as Negative + Negative regulation predicted as Negative + Regulation of transcription predicted as Negative + Regulation of gene expression predicted as Negative + Catalysis of phosphorylation predicted as Negative + Catalysis of posttranslational modification predicted as Negative + Catalysis of ubiquitination predicted as Negative + Catalysis of methylation predicted as Negative + Catalysis of dephosphorylation predicted as Negative + Catalysis of acetylation predicted as Negative + Regulation of degradation predicted as Negative + Catalysis of SUMOylation predicted as Negative + Regulation predicted as Complex formation + Catalysis of deneddylation predicted as Negative + Catalysis of glycosylation predicted as Negative + Catalysis of posttranslational modification predicted as Complex formation + Catalysis of palmitoylation predicted as Negative + Negative regulation predicted as Complex Formation + Catalysis of ADP-ribosylation predicted as Negative + Catalysis of deacetylation predicted as Negative + Catalysis of deubiquitination predicted as Negative + Catalysis of other small molecule conjugation or removal predicted as Negative + Catalysis of small protein conjugation or removal predicted as Negative + Regulation of gene expression predicted as Complex formation + Catalysis of demethylation predicted as Negative + Catalysis of neddylation predicted as Complex formation + Catalysis of phosphoryl group conjugation or removal predicted as Negative + Catalysis of small protein conjugation predicted as Negative + Other catalysis of small protein conjugation predicted as Negative + Positive regulation predicted as Complex formation + Regulation of degradation predicted as Complex formation = 201 + 137 + 128 + 52 + 38 + 30 + 21 + 17 + 15 + 13 + 10 + 9 + 7 + 7 + 6 + 6 + 6 + 5 + 5 + 4 + 4 + 3 + 2 + 2 + 2 + 1 + 1 + 1 + 1 + 1 + 1 + 1 = 737

##### *Assigned a directed interaction, where there should be none*

Negative predicted as Regulation + Negative predicted as Positive Regulation + Negative predicted as Negative Regulation + Negative predicted as Regulation of transcription + Negative predicted as Catalysis of ubiquitination + Negative predicted as Regulation of degradation + Negative predicted as Catalysis of phosphorylation + Negative predicted as Catalysis of methylation + Negative predicted as Regulation of gene expression + Negative predicted as Catalysis of dephosphorylation + Negative predicted as Catalysis of acetylation + Complex formation predicted as Regulation + Negative predicted as Catalysis of posttranslational modification + Complex formation predicted as Negative regulation + Complex Formation predicted as Positive Regulation + Negative predicted as Catalysis of deacetylation + Negative predicted as Catalysis of palmitoylation + Negative predicted as Catalysis of demethylation + Negative predicted as Catalysis of ADP-ribosylation + Negative predicted as Catalysis of glycosylation + Negative predicted as Catalysis of SUMOylation = 156 + 130 + 113 + 44 + 34 + 33 + 15 + 14 + 16 + 12 + 11 + 10 + 7 + 4 + 4 + 4 + 4 + 3 + 2 + 2 + 2 = 620

##### *Assigned a directed interaction, but the direction is wrong*

Regulation predicted as Regulation (reverse) + Negative Regulation predicted as Negative Regulation (reverse) = 3+2 = 5

##### *Correctly assigned a sign*

$TP_{\text{Negative regulation}} + TP_{\text{Positive regulation}} = 276 + 263 = 539$

*Failed to assign a sign (positive or negative), where there should be one*

Positive regulation predicted as Regulation + Negative regulation predicted as Regulation + Negative regulation predicted as Regulation of gene expression + Negative regulation predicted as Regulation of transcription + Positive regulation predicted as Catalysis of palmitoylation = 19 + 9 + 1 + 1 + 1 = 31

*Assigned a sign, where there should be none*

Regulation predicted as Positive Regulation + Regulation predicted as Negative Regulation + Regulation of degradation predicted as Negative Regulation + Catalysis of acetylation predicted as Positive regulation + Catalysis of demethylation predicted as Positive regulation + Catalysis of glycosylation predicted as Negative Regulation + Catalysis of methylation predicted as Negative regulation + Regulation of transcription predicted as Positive Regulation = 26 + 12 + 3 + 2 + 1 + 1 + 1 + 1 = 47

*Assigned a sign, but the sign is wrong*

Negative Regulation predicted as Positive Regulation + Positive Regulation predicted as Negative Regulation = 7 + 7 = 14
